# Comprehensive multi-omics single-cell data integration reveals greater heterogeneity in the human immune system

**DOI:** 10.1101/2021.07.25.453651

**Authors:** Congmin Xu, Junkai Yang, Astrid Kosters, Benjamin R. Babcock, Peng Qiu, Eliver E. B. Ghosn

**Author notes:** Contributed equally (Congmin Xu, Junkai Yang). Corresponding authors: Eliver E.B. Ghosn | Peng Qiu |.

## Abstract

Single-cell transcriptomics enables the definition of diverse human immune cell types across multiple tissues and disease contexts. Still, deeper biological understanding requires comprehensive integration of multiple single-cell omics (transcriptomic, proteomic, and cell-receptor repertoire). To improve the identification of diverse cell types and the accuracy of cell-type classification in multi-omics single-cell datasets, we developed SuPERR-seq, a novel analysis workflow to increase the resolution and accuracy of clustering and allow for the discovery of previously hidden cell subsets. In addition, SuPERR-seq accurately removes cell doublets and prevents widespread cell-type misclassification by incorporating information from cell-surface proteins and immunoglobulin transcript counts. This approach uniquely improves the identification of heterogeneous cell types in the human immune system, including a novel subset of antibody-secreting cells in the bone marrow.

## Introduction

Single-cell RNA sequencing (scRNA-seq) technologies have rapidly advanced in the last decade, including advances to cell-capture approaches (Evan et al. 2015; Allon et al. 2015; Utada et al. 2007), library preparation (Picelli et al. 2013; Hashimshony et al. 2012), and sequencing methods (Evan et al. 2015; Picelli et al. 2013; Habib et al. 2017; Stoeckius et al. 2017). These increasingly more widely adopted technologies have significantly improved the understanding of cell heterogeneity in health and disease (Hashimshony et al. 2012; Zheng et al. 2017; Habib et al. 2017; Stoeckius et al. 2017; Picelli et al. 2013). However, reliance on cellular transcriptomics alone limits the comprehensive identification of heterogenous cell populations (Liu and Trapnell 2016). This limitation has propelled the development of multi-omics single-cell sequencing technologies to increase the resolution and accuracy for cell subset classification.

Multi-omics single-cell sequencing technologies, such as CITE-seq (Stoeckius et al. 2017), REAP-seq (Peterson et al. 2017), and others (Lee, Hyeon, and Hwang 2020), simultaneously measure gene expression (mRNA) and cell-surface proteins. Additional heterogeneity of immune cell subsets can be revealed by combining single-cell gene expression with simultaneous T- and B-cell receptor (TCR and BCR) repertoire sequencing using techniques such as RAGE-seq and DART-seq (Meyer 2019; Singh et al. 2019; Horns, Dekker, and Quake 2020; Zemmour et al. 2018; Yermanos et al. 2021). Thus, simultaneous measurement and comprehensive integration of transcriptomics, cell-surface protein, and cell-receptor repertoire can reveal heterogeneous cell types relevant to disease mechanisms and homeostasis.

However, multi-omics technologies also present computational challenges for data integration and analysis (Colomé-Tatché and Theis 2018; Luecken and Theis 2019; Stuart and Satija 2019). Challenges include high dimensionality of the data (Yu and Lin 2016), sparsity of the data (Qiu 2020), diversity across various omics data types (Hao et al. 2021), and technical effects between different sample batches (Stuart et al. 2019). Several algorithms have been developed to integrate and analyze multi-omics measurements, including weighted nearest neighbor (WNN) implemented in Seurat v4 (Hao et al. 2021), similarity network fusion (SNF) in CiteFuse (Kim et al. 2020), among others (Wang et al. 2020; Gayoso et al. 2021; Jin, Zhang, and Nie 2020; Argelaguet et al. 2018). The commonality of these methods is to utilize the shared signals among different omics data types to align their distributions and achieve integration, which is an unsupervised data-driven approach. While unsupervised data-driven methods have been successful for clustering and identifying cell types, significant improvements can be made by incorporating robust prior knowledge such as well-established marker genes and cell-surface protein markers that can accurately define cell types (Aran et al. 2019; Mahnke, Chattopadhyay, and Roederer 2010).

Here, to address the challenges of multi-omics analysis, we combined our extensive expertise on high-dimensional flow cytometry data analysis (Meehan et al. 2019) with our multi-omics single-cell data sets to develop the SuPERR-seq (**<u>Su</u>**rface **<u>P</u>**rotein **<u>E</u>**xpression, m**<u>R</u>**NA- and **<u>R</u>**epertoire-seq) workflow. SuPERR-seq is a novel, semi-supervised, biologically-motivated approach towards the integration and analysis of multi-omics single-cell data matrices. By combining a robust prior knowledge of flow cytometry-based cell-surface markers (gating strategy) (Mahnke, Chattopadhyay, and Roederer 2010) with the high-dimensional analysis of scRNA-seq, SuPERR-seq increases the resolution and accuracy in clustering algorithms and allows the discovery of new biologically relevant cell subsets. We first applied the flow cytometry-based “gating strategy” on a combination of cell-surface markers and immunoglobulin-specific transcript counts to identify major immune cell lineages. Next, we explored the gene expression matrix following this gating strategy to resolve cell subsets within each major immune lineage. The inclusion of this atypical “gating strategy” step also allows for cell-doublet discrimination and dramatically enhances lineage-specific variation, which helps better capture biological signals *within* each cell lineage. Finally, we apply the SuPERR-seq workflow to human blood and bone marrow cells and directly compare its performance to existing methods. We show that SuPERR-seq can leverage the power of each “omics” to identify major immune lineages more accurately and reveal biologically-meaningful heterogeneity within each lineage that can be confirmed by flow cytometry, facilitating the discovery of novel immune cell types.

## Results

### Cell-surface proteins and immunoglobulin transcript counts identify major immune lineages in human blood and bone marrow

We generated multi-omics single-cell datasets from 12,759 human peripheral blood mononuclear cells (PBMC) and 7,426 human bone marrow (BM) cells from five healthy adult donors. As shown in Fig. 1A, we simultaneously captured the following three omics: total gene expression (GEX), 32 cell-surface proteins/antibody-derived tags (ADT) (Table S1), and B-cell receptor (BCR) heavy and light chain V(D)J repertoire (VDJ). The SuPERR-seq analysis workflow can be described in two major steps shown in Fig. 1B: 1) a manual biaxial gating based on the expression levels of well-stablished(Mahnke, Chattopadhyay, and Roederer 2010) cell-surface proteins (ADT) and total immunoglobulin (Ig) transcript counts to accurately identify antibody-secreting cells, and 2) a subsequent sub-clustering of each manually-gated lineage/population identified in step 1, using the GEX matrix.

**Figure 1.**
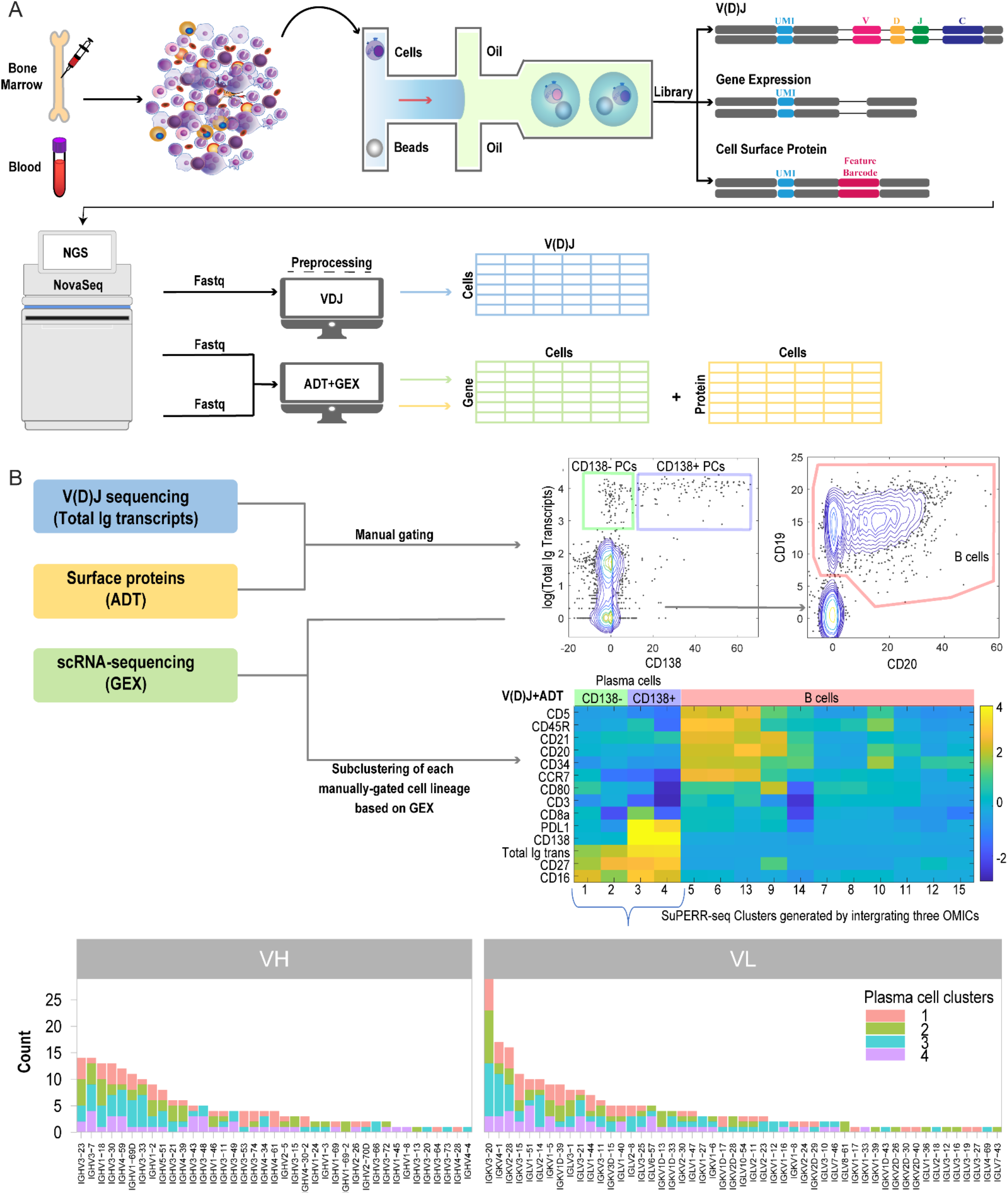
SuPERR-seq workflow. (A) Schematic overview of the experimental design. Peripheral blood and bone marrow aspirates were processed, surface-stained with barcoded antibodies, and then encapsulated with barcoded microspheres. We generated three libraries for each sample corresponding to gene expression (GEX), cell-surface protein/antibody-derived tags (ADT), and cell-receptor repertoire (VDJ). Libraries were sequenced to a target depth, and count matrices were assembled for each-omic data separately. (B) SuPERR-seq workflow is composed of two main steps. Major cell lineages are manually gated at the first step by integrating information from both the ADT and V(D)J data matrices. Then, the manually-gated cell lineages are further sub-clustered based on information from the GEX data. The V(D)J matrix can be used to further identify the diversity of heavy (VH) and light (VL) variable genes among the plasma cell clusters. PCs: plasma cells.

For the first step of the SuPERR-seq workflow, we normalized the cell-surface protein (ADT) data using the DSB normalization method (Mulè, Martins, and Tsang 2021). Next, we concatenated the normalized ADT matrix with the total Ig-specific unique molecular identifier (UMI) counts from the V(D)J matrix, which describes the total number of immunoglobulin-derived transcripts per cell. The integrated ADT/Ig matrix was used to identify major immune cell lineages before assessing their gene expression profile. Major immune cell lineages were identified and classified using a well-established sequential gating strategy on biaxial plots (Figs. 2A and 3A) widely used in conventional flow cytometry data analysis and readily available through the Optimized Multicolor Immunofluorescence Panel (OMIP) publications (Mahnke, Chattopadhyay, and Roederer 2010). Since antibody-secreting cells (ASC), also known as plasma cells, produce and secrete large quantities of immunoglobulin, they could be accurately identified based on their Ig-specific transcript counts (Figs. 2A and 3A). Of note, the semi-supervised SuPERR-seq workflow was able to readily identify a rare cell cluster containing as few as eight plasma cells in the human PBMC sample. As we show below, such a rare population of plasma cells cannot be identified using current conventional unsupervised workflows.

**Figure 2.**
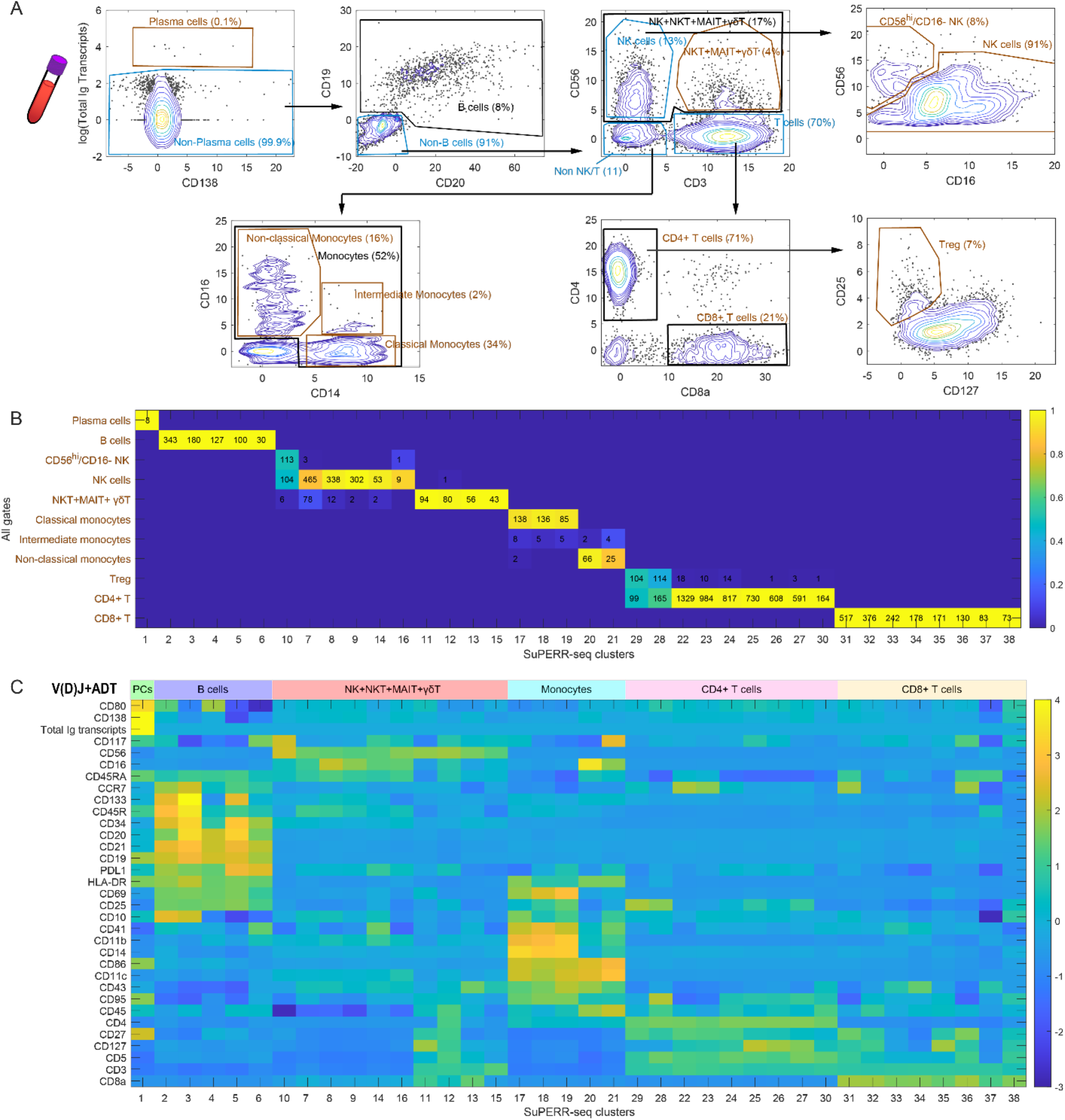
SuPERR-seq workflow applied to PBMCs. (A) “Gating strategy” approach to identify major cell lineages on biaxial plots based on surface markers (ADT) and V(D)J data. Gates for major lineages are indicated as black outlines and black text. Gates for downstream cell-identity validation are indicated as golden outlines and golden text. (B) Cross comparison between the manually-gated major lineages and the final SuPERR-seq clusters. (C) The average expression levels of surface markers (ADT) and Ig-specific transcripts (VDJ) for the final SuPERR-seq clusters. PCs: plasma cells.

**Figure 3.**
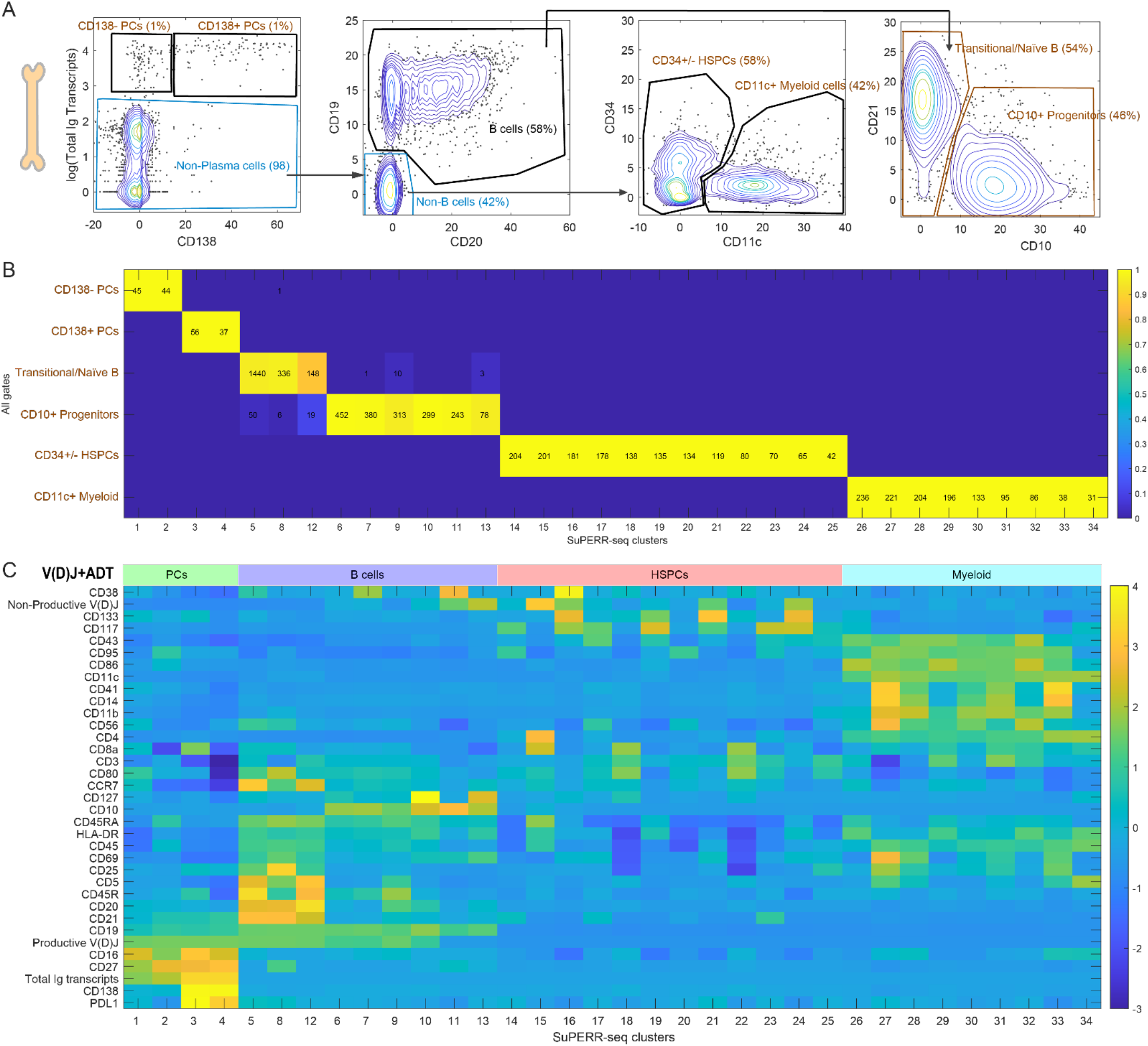
SuPERR-seq workflow applied to BM cells. (A) “Gating strategy” approach to identify major cell lineages on biaxial plots based on surface markers (ADT) and V(D)J data. Gates for major lineages are indicated as black outlines and black text. Gates for downstream cell-identity validation are indicated as golden outlines and golden text. (B) Cross comparison between the manually-gated major lineages and the final SuPERR-seq clusters. (C) The average expression levels of surface markers (ADT) and Ig-specific transcripts (VDJ) for the final SuPERR-seq clusters. PCs: plasma cells.

In the analysis of PBMC samples, we defined gates for six major immune cell lineages based on well-established markers (see Table 1): plasma cells, B cells, NK/NKT/MAIT/γδT cells, Monocytes, CD4+ T cells, and CD8+ T cells (Fig. 2A, black borders). In the analysis of the BM samples, we defined gates for five major lineages (see Table 2): CD138-Plasma cells, CD138+ Plasma cells, B cells, Myeloid cells, and Hematopoietic Stem and Progenitor Cells (HSPCs) (Fig. 3A, black borders). Notably, the B cells identified by the manual gating strategy using the cell-surface markers (ADT) were also present within the V(D)J matrix, which validated our strategy of using ADT for cell-lineage classification (Figs. S1, S2, S3A). In addition to the main cell lineages, our manual-gating strategy revealed other sub-clusters (Figs. 2A and 3A, gray borders), which we used to validate the results from the downstream GEX-based clustering analysis. Our manual-gating strategy was further validated by high-dimensional flow cytometry analysis using an aliquot of the same samples taken before the single-cell encapsulation (Fig. S3B).

**Table 1.**
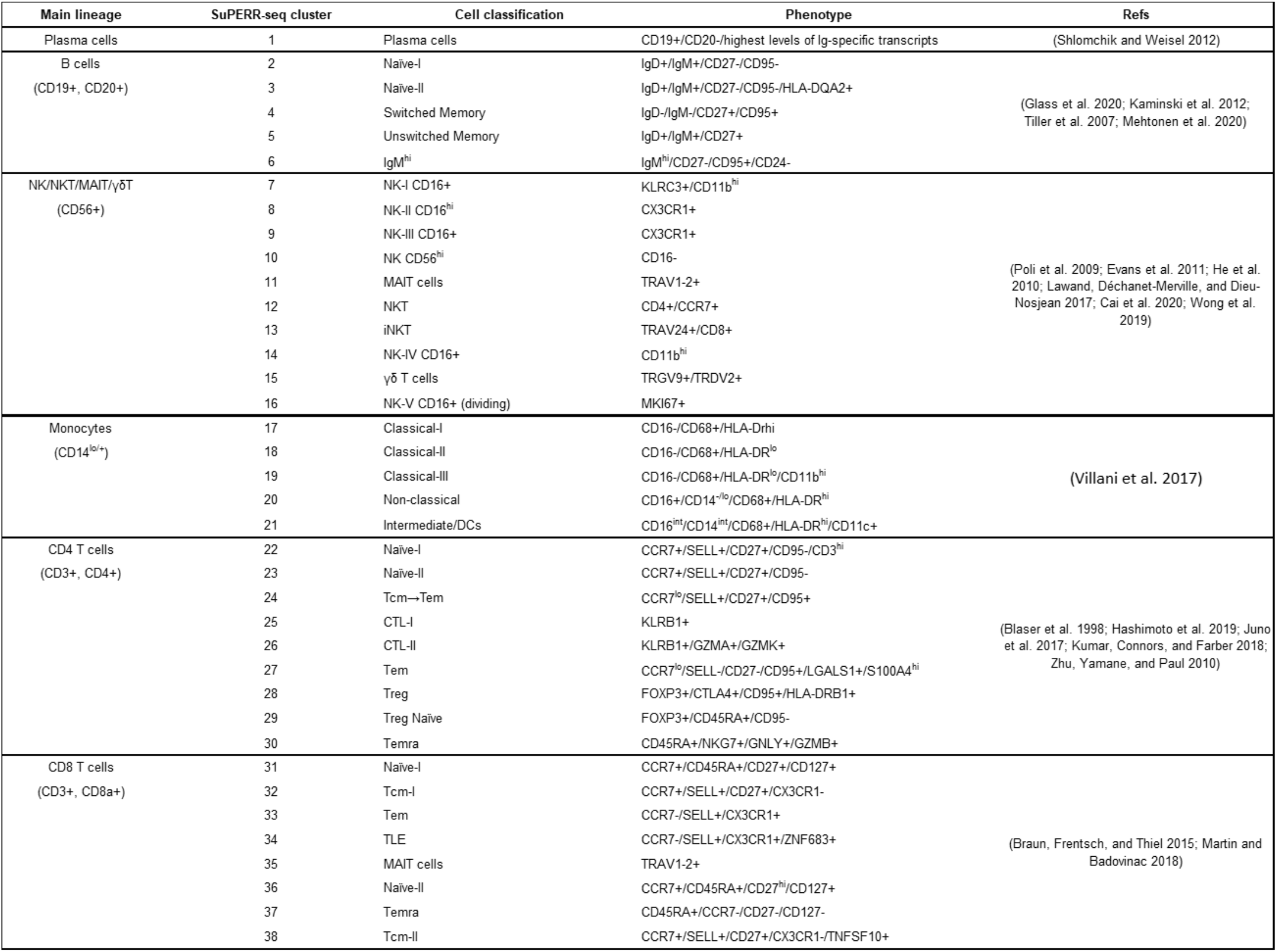
Cell-type classification and phenotype of PBMC clusters generated by the SuPERR-seq workflow. Gene expression (GEX) and cell-surface (ADT) markers used to classify the lymphoid and myeloid cell populations in the human peripheral blood mononuclear cells (PBMC). NK: Nature Killer cells; NKT: NK T cells; iNKT: invariant NKT; γδ T: gamma-delta T cells; Tcm: central memory T cells; Tem: effector memory T cells; CTL: cytotoxic T lymphocytes; Treg: regulatory T cells; Temra: effector memory T cells expressing CD45RA; TLE: long-lived effector memory T cells; MAIT: mucosal-associated invariant T cells.

**Table 2.**
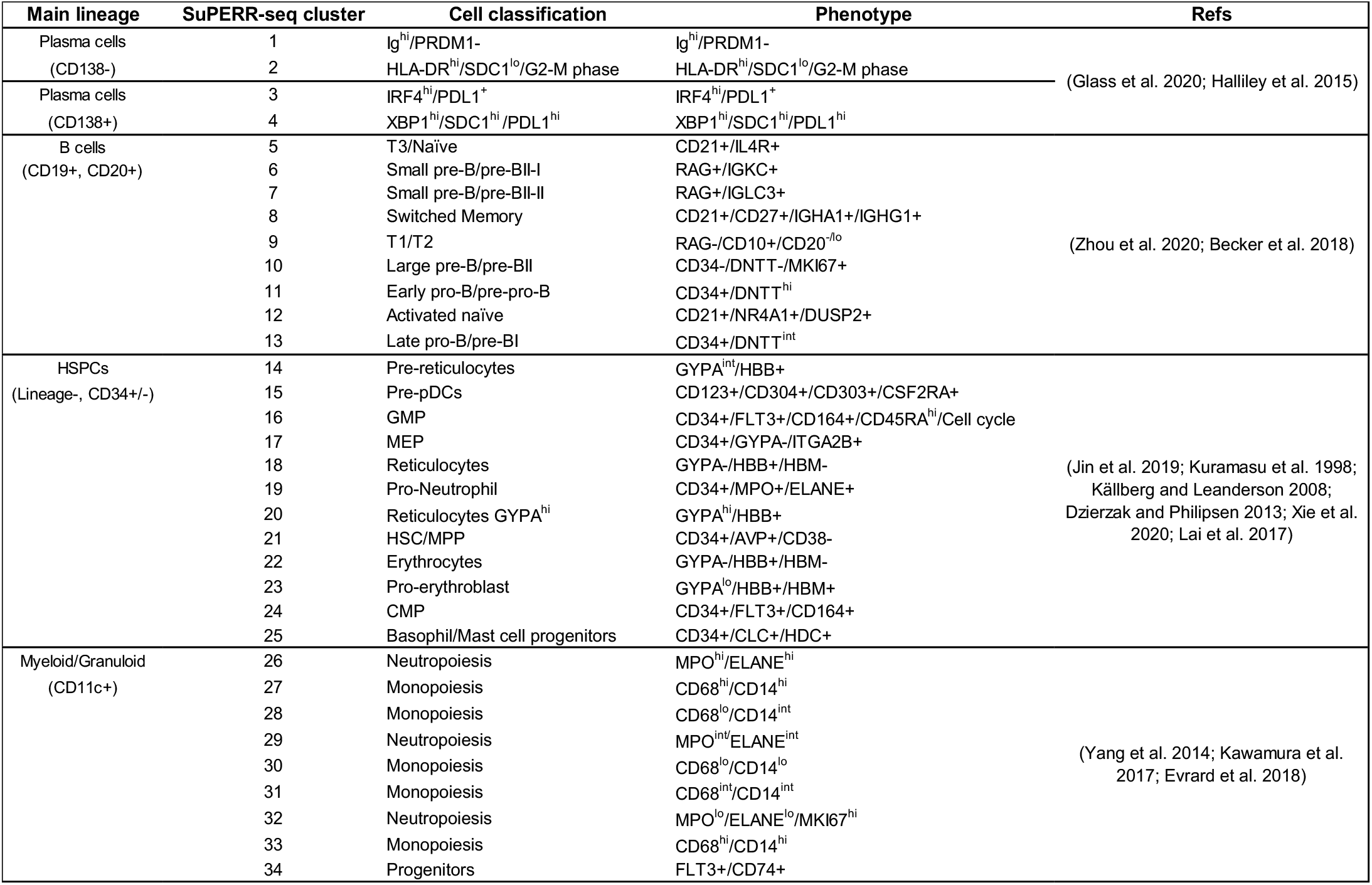
Cell-type classification and phenotype of BM clusters generated by the SuPERR-seq workflow. Gene expression (GEX) and cell-surface (ADT) markers used to classify the plasma cells, B cells, myeloid and granuloid cells, and the hematopoietic stem and progenitor cells (HSPCs) in the human bone marrow (BM). Ig: immunoglobulin transcripts; G2-M phase: genes involved in the cell cycle; T1/2/3: Transitional B cells; pDCs: plasmacytoid dendritic cells; GMP: granulocyte-monocyte progenitor; MEP: megakaryocyte-erythroid progenitor; HSC: hematopoietic stem cell; MPP: multipotent progenitor; CMP: common myeloid progenitor.

#### Cell-doublet discrimination

By identifying the *major cell lineages* as the first step of the SuPERR-seq workflow we were able to accurately identify and remove cell doublets. For example, we applied flow cytometry-style biaxial feature plots on the ADT data to identify cell barcodes that co-expressed features of two or more major lineages (Figs. S1 and S2). Cell barcodes containing ADT signals (cell-surface markers) known to belong to more than one well-defined immune lineage (e.g., co-expression of B-cell/CD19 and T-cell/CD3 markers) were considered doublets and removed from the downstream analysis. Figs. S1 and S2 illustrate the gating strategy used for removing doublets from the PBMC and BM data sets, respectively. In our benchmarking analysis below, we further validate and compare our cell-doublet discrimination method here to other recently-developed algorithms.

#### Selection of major cell lineages prior to GEX analysis enhances true biological signals

Notably, the assignment of major cell lineages via a supervised manual-gating approach (ADT + V(D)J matrices) before performing the principal component analysis (PCA) on the GEX matrix, revealed that much of the data variation captured by the PCA is cell-lineage specific (Fig. 4). Key variables, including ribosomal transcript content, mRNA abundance, and total unique gene counts, vary significantly among the cell lineages (Fig. 4, ANOVA p<2.2e-16). Even though our initial step of sample integration only outputted data for 2000 highly variable genes (HVGs), meaning that the subsequent PCA for the various lineages were performed on the same set of HVGs, the PCA can be interpreted as a feature selection step. For example, the PCA analysis focusing only on a pre-defined subset of cells (i.e., pre-gated major lineages) is able to produce principal components driven by biologically-meaningful variations that occur only *within* that lineage. In contrast, the principal components computed based on all cells are primarily driven by variations among different lineages. As such, enhancing the lineage-specific biological signals should also capture variations originated from differences in cellular states within a particular major lineage. Thus, lineage-specific variations, including variations due to differences in cellular state, will be reflected in the corresponding PCA, which in turn will be reflected in the downstream clustering and visualization (UMAP/t-SNE) results for the corresponding lineage. In sum, our pre-selection of major cell lineages before GEX clustering generates lineage-specific principal components that are biologically-meaningful, improving the resolution of the downstream GEX clustering analysis.

**Figure 4.**
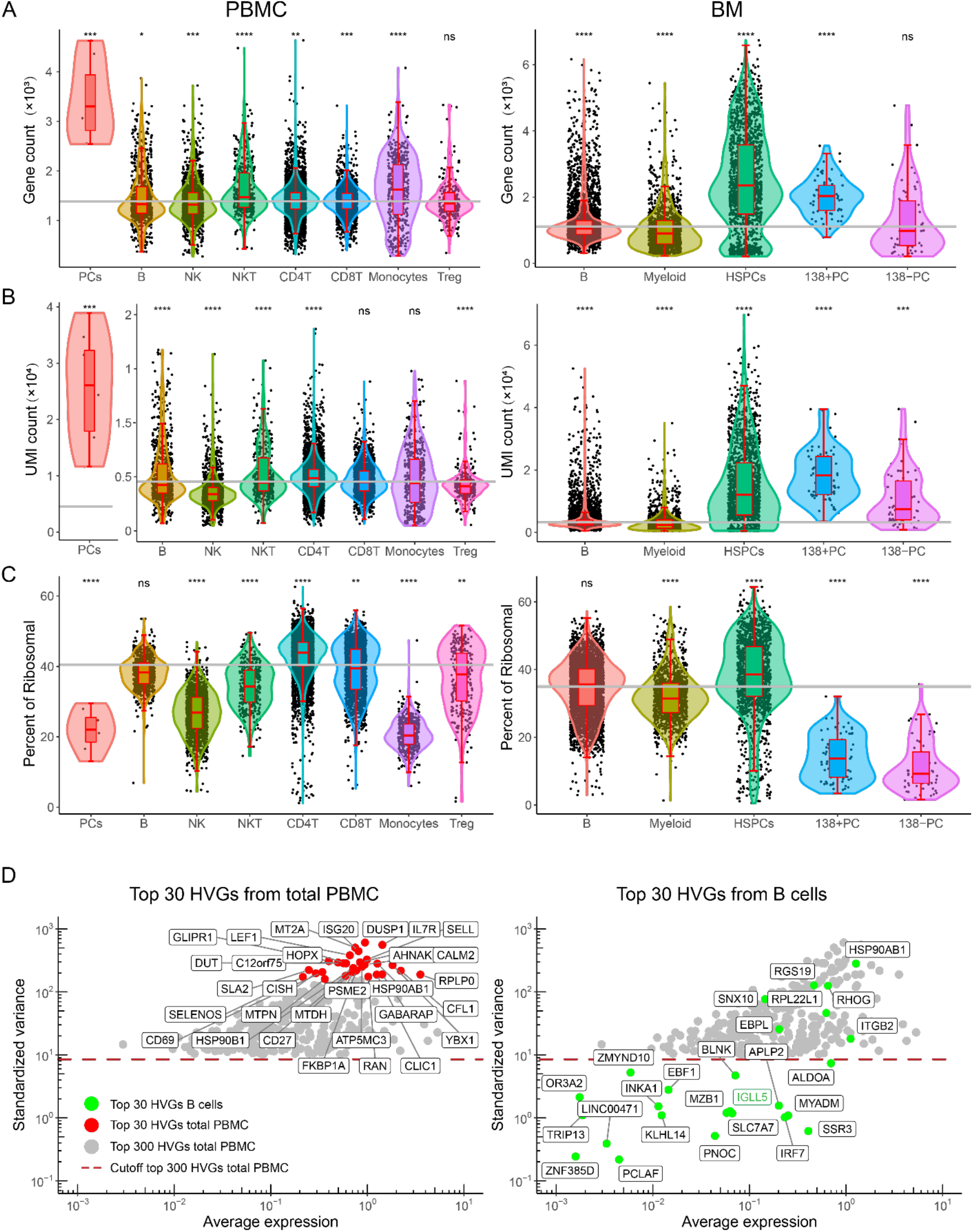
Cell-type-specific variations in gene expression. (A) “Gene count” represents the number of unique genes expressed by each cell type. (B) “UMI count” represents the total mRNA abundance expressed by each cell type. (C) “Percent of Ribosomal” represents the percentage of ribosomal gene UMI counts expressed by each cell type. The grey line shows the mean expression level for each feature in the total PBMC and BM samples. (D) Left panel: the top 30 (red points) and the top 300 (grey points) HVGs from total PBMC. The points under the red dashed line fall below the top 300 HVGs of total PBMC. Right panel: the top 30 highly-variable genes (HVGs) from PBMC-derived B cells (green points) displayed with the top 300 HVGs from total PBMC (grey points). Student’s t-test was used to compare the mean of each cell type with the mean of the total PBMC/BM. *p<0.05, **p<0.01, ***p<0.001, ****p<0.0001, unpaired, two-tailed. Multiple-group ANOVA test for (A), (B), and (C): p<2.2e-16. PCs: plasma cells.

### SuPERR-seq reveals greater heterogeneity within major immune lineages

#### Human Peripheral Blood

Gating for major cell lineages using the ADT and V(D)J matrices revealed six distinct populations within the PBMC sample (Fig. 2A, black outlines). We then further explored each major lineage by generating subclusters using information from the GEX matrix. Briefly, we selected a set of HVGs from within the pre-gated population. The counts of selected genes for each cell were normalized by library size and then natural-log transformed, followed by per-gene Z-score scaling. We then applied a singular value decomposition (SVD) implementation of principal component analysis (PCA) to reduce the dimensionality. Left singular values were taken as gene scores and right singular values as cell scores. Next, we generated a K-nearest neighbors’ graph, followed by Louvain community detection (see Methods section for detailed description). Following these steps, we obtained the subclusters for each major lineage, herein called SuPERR-seq clusters. At a Louvain resolution of 0.8, SuPERR-seq identified 38 clusters in human PBMC, with each major lineage broken out into two or more subclusters, representing sub-lineage heterogeneity (Fig. 2B).

We further investigated the 38 SuPERR-seq clusters by exploring the expression levels of cell-surface proteins (ADT matrix) (Figs. 2C and S5). Some ADT markers were lineage-specific (e.g., CD19 was used to classify B cells and Ig-specific transcript counts were used to classify plasma cells) and used to confirm SuPERR-seq cell classification accuracy. Other markers displayed heterogeneous expression within lineages and were primarily used for defining and confirming subcluster identity. We integrated the ADT and GEX data matrices by simple concatenation to generate a joint matrix for SuPERR-seq cluster identification (Figs. 5 and S4-S11). We confirmed the heterogenous clusters as biologically meaningful using well-established cell-surface lineage markers (Table 1). For example, five subclusters were identified within the B-cell major lineage, all of which could be mapped back to previously described B-cell subsets (Garimalla et al. 2019; Glass et al. 2020; Kaminski et al. 2012; Tiller et al. 2007) (Fig. S5). Similarly, several subclusters were identified within the T-cell and NK-cell major lineages (Fig. S7-S9), including T-regulatory cells (Treg) defined as a subtype of CD4+ T cells with high expression of surface CD25 (ADT matrix) and FOXP3 transcripts (GEX matrix), and low expression of surface CD127 (ADT matrix) (Figs. 2A and S7). Finally, five subclusters of monocytes were identified, including the previously described classical, non-classical, and intermediate monocyte subsets (Fig. S6).

**Figure 5.**
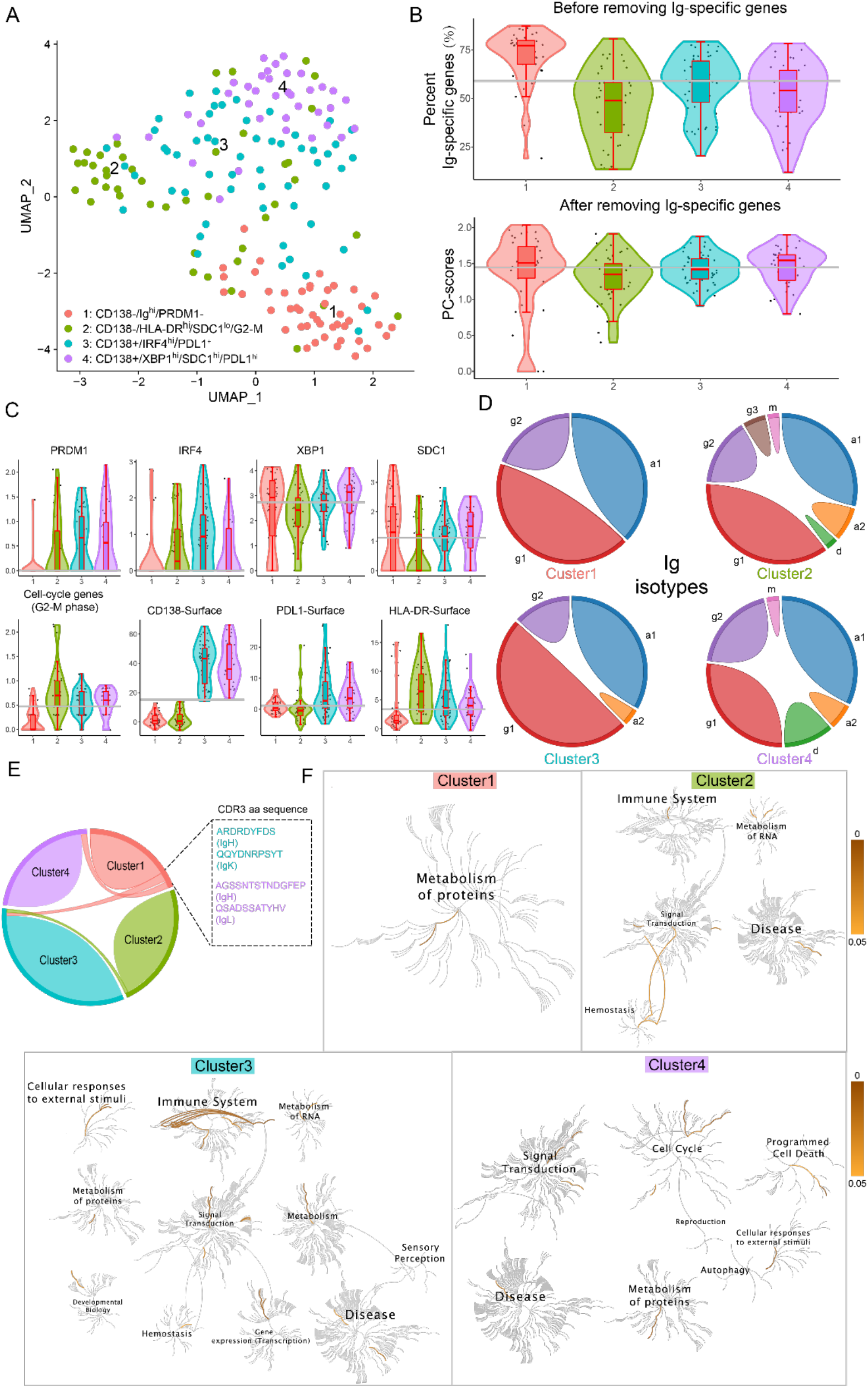
SuPERR-seq workflow identifies four subsets of human plasma cells in the BM. (A) UMAP representation of the four bone marrow (BM) plasma cells. (B) Top panel: percentage of Ig-specific transcripts (UMI) expressed in each plasma cell subset. Bottom panel: expression levels (sum) of plasma cell genes (see Methods) after removing Ig-specific UMIs and re-normalizing the data matrix. (C) Expression levels of individual plasma cell genes, cell-cycle score after removing Ig-specific UMIs (See Methods) and ADT. The grey line shows the mean expression level across all clusters. (D) The antibody isotypes and subclasses expressed by each plasma cell subset. (E) The connected lines on the Circus plot describe shared clones between clusters (clonal lineage was identified by the identical V and J gene usage, identical CDR3 nucleotide length, and ≥85% homology within the CDR3 nucleotide sequence). (F) Reactome Pathway Database analysis (see Methods) shows unique biological processes that define each plasma cell subset.

Remarkably, SuPERR-seq readily identified a cluster of rare plasma cells (antibody-secreting cells) in the blood even though we captured only eight plasma cells within this cluster (Fig. 2). This level of resolution and accuracy in identifying rare plasma cells was possible by analyzing the total Ig-specific transcript counts from the V(D)J matrix, which unlike the analogous Ig transcripts from the GEX matrix, provide a more accurate count of the total productive Ig expression per cell. As plasma cells are defined by their unique ability to produce large quantities of Ig transcripts, they were readily identified based on a ∼2.5 log_10_-fold increase in total Ig-specific UMI counts compared to B cells (Fig. 2A).

#### Human Bone Marrow

SuPERR-seq identified 34 unique clusters in human BM cells (Figs. 3A and 3B). Of note, SuPERR-seq matched the various B-cell subclusters to the different stages of B-cell development known to occur in the human BM (Mehtonen et al. 2020) (Fig. S4). For example, the cell-surface expression of CD10, along with DNTT and CD34 gene expression (GEX) transcripts, classified cluster 11 and cluster 13 as Early Pro-B (a.k.a., pre-pro-B) and Late Pro-B (a.k.a., pre-BI), respectively. The lack of DNTT mRNA transcripts in cluster 10 indicates a Large Pre-B stage (a.k.a., pre-BII), which is followed by the Small Pre-B stage (a.k.a., pre-BII) represented by clusters 6 (VPREB1^hi^) and 7 (VPREB1^lo^). The Transitional (T)1/T2/T3 and Naive B cells could be identified by their surface expression of CD21 (Zhou et al. 2020). Finally, the mature class-switched memory B cells were identified based on their expression of IGHA1/IGHG1 immunoglobulin transcripts and cell-surface CD27 (Becker et al. 2018) (Fig. S4). Similarly, the SuPERR-seq workflow also identified the developmental pathway of neutrophils, monocytes, and erythrocytes starting from the most undifferentiated population of hematopoietic stem cells (HSC) and multipotent progenitors (MPP) expressing CD34 and AVP transcripts (but lacking CD38) (Figs. S10 and S11). The classification results for the BM subclusters are summarized in Table 2.

### SuPERR-seq workflow reveals new subsets of antibody-secreting cells in the human bone marrow

The SuPERR-seq workflow readily and unambiguously identified four biologically distinct subsets of human antibody-secreting cells (a.k.a., plasma cells) in the adult bone marrow (BM) of healthy donors (Figs. 3 and 5). The determinant feature of a plasma cell is its ability to produce and secrete large quantities of immunoglobulins (Ig) (i.e., antibodies). The SuPERR-seq workflow leverages this unique plasma cell feature by quantifying the absolute UMI counts of Ig-specific genes (IGH + IGL) from the V(D)J repertoire matrix (Fig. 3A). Next, we integrate the Ig UMI count matrix to the ADT matrix we used to pre-gate major lineages (see above) and apply the same semi-supervised gating strategy on biaxial plots to identify cells with high Ig UMI counts. Unlike the GEX matrix, the Ig-specific UMI counts from V(D)J matrix provides a more accurate count of the productive Ig transcripts produced by plasma cells as the V(D)J data matrix is generated from a separate library using only Ig-specific primers (Zheng et al. 2017).

Remarkably, the number of total Ig-specific transcripts (UMI counts) detected in plasma cells is ∼2.5 log_10_-fold higher than in B cells (Figs. 3A and 3C), allowing for an unambiguous identification of total plasma cells. Next, we separated the total plasma cells into two subsets, based on the cell-surface expression of CD138 (ADT matrix), a canonical plasma-cell marker expressed by some, but not all, BM plasma cells (Halliley et al. 2015). Finally, we used the GEX matrix to further subdivide the CD138+ and CD138-plasma cell subsets based on their transcriptomic profile, revealing a total of 4 distinct subsets of human plasma cells (Figs. 3 and 5).

To facilitate the identification of the differential gene expression that distinguish each of the four plasma cell subsets, we first removed from the GEX matrix all the mRNA transcripts derived from immunoglobulin genes (i.e., we removed IGHV, IGKV and IGLV genes). The rationale is that immunoglobulin genes represent more than 50% of the total mRNA transcripts (UMI counts) recovered from plasma cells. We then log-normalized the immunoglobulin-depleted GEX matrix and performed DGE analysis using the Wilcoxon Rank-Sum Test followed by Bonferroni correction. The resulting differentially-expressed genes readily defined unique biological processes for each plasma cell cluster (Fig. 5). Cluster 1: CD138-/Ig^hi^/PRDM1-; cluster 2: CD138-/HLA-DR^hi^/SDC1^lo^/G2-M phase; cluster 3: CD138+/IRF4-/PDL1+, cluster 4: CD138+/XBP1^hi^/SDC1^hi^/PDL1^hi^ (Figs. 5A and 5C). Clusters 2 and 4 show characteristics similar to previously identified human BM plasma cell subsets, described as Fraction A and Fraction B, respectively (Halliley et al. 2015). Notably, cluster 1 represents a unique plasma cell subset, in that >75% of its total mRNA transcripts represent immunoglobulin genes (Fig. 5B). This large proportion of immunoglobulin gene transcripts indicates a high metabolic activity that is geared towards producing and secreting antibodies. Indeed, pathway analysis (Reactome Pathway Database) (Jassal et al. 2020) using the DGE list for cluster 1 revealed signals mainly for the metabolism of proteins (Fig. 5F).

Surprisingly, we found that not all plasma cell clusters express the canonical plasma cell genes. Plasma cells develop from activated B cells through a dynamic cell-differentiation process, leading to the down-regulation of B-cell identity genes, such as PAX5, and up-regulation of well-described plasma-cell genes, including PRDM1, SDC1, XBP1, and IRF4 (Halliley et al. 2015). These genes are considered canonical plasma-cell genes and, hence, they are used in scRNA-seq experiments to identify and classify plasma cells based on their transcriptomics (GEX matrix). However, the plasma cell cluster 1 does not express detectable levels of IRF4 and PRDM1. The absence of these canonical genes in cluster 1 was not due to the overrepresentation of immunoglobulin (Ig) genes because IRF4 and PRDM1 were not detected even after normalizing the GEX matrix without the immunoglobulin genes (Fig. 5C).

By comparing the immunoglobulin isotypes and subclasses (IgM, IgD, IgG1-4, IgA1-2, IgE) expressed by each plasma-cell subset identified by SuPERR-seq, we found that cluster 2 contains plasma cells of multiple isotypes/subclasses. In contrast, cluster 1 is more homogeneous, containing mainly IgG1 and IgA1 (Fig. 5D). Furthermore, we observed that cluster 1 is composed of clonal plasma cells (defined by their IGH CDR3 amino acid sequences) that are shared among clusters 3 and 4 (Fig. 5E). Finally, pathway analysis (Reactome Pathway Database) revealed unique biological processes and genetic programs that define each plasma cell subset (Fig. 5F). Notably, cluster 4 expresses genes involved in cell cycle and programmed cell death, while cluster 3 appears to be actively responding to environmental stimulation.

In sum, the SuPERR-seq workflow readily and unambiguously identified four biologically-distinct human plasma cell subsets in the adult BM. These findings further support the need for comprehensive multi-omics single-cell data integration and reveal the potential shortcomings of relying solely on one omics data type (i.e., transcriptomics) to identify and classify cell (sub)types. In the following sections, we provide specific examples in which SuPERR-seq workflow can outperform existing approaches.

### Benchmarking of SuPERR-seq against current methods developed to remove cell doublets

As we described above, the first step of the SuPERR-seq workflow, in which we use cell-surface markers (ADT) and V(D)J gene counts in sequential biaxial plots to identify major cell lineages, also allows for accurate cell-doublet discrimination (Figs. S1, S2, 6A, and 6B). Our SuPERR-seq approach successfully identified and removed 370 cell doublets in the PBMC sample. In contrast, standard/conventional approaches (Ocasio et al. 2019) of trimming out cell barcodes with very high mRNA transcripts (e.g., removing cell barcodes with greater than the mean UMI value + 4 standard deviation) identified only 42 doublets. In the BM, SuPERR-seq identified 108 cell doublets, and the conventional approach identified only one cell doublet (Fig. 6A). Strikingly, every plasma cell we identified in PBMCs would have been trimmed/removed by the conventional approach of removing cells with very high mRNA transcripts (i.e., mean + 4 SD UMI counts) (Fig. 6A). In contrast, the SuPERR-seq workflow readily recognized the PBMC plasma cells as single cells containing high UMI counts (Fig. 6A). The plasma cell identity was further confirmed by the presence of a single productive V(D)J repertoire usage and the lack of other major lineage markers. Thus, the true cell doublets removed by SuPERR-seq would otherwise be missed by conventional approaches and erroneously included as single cells in downstream GEX analysis, as visualized on the UMAP plot (Fig. 6C).

**Figure 6.**
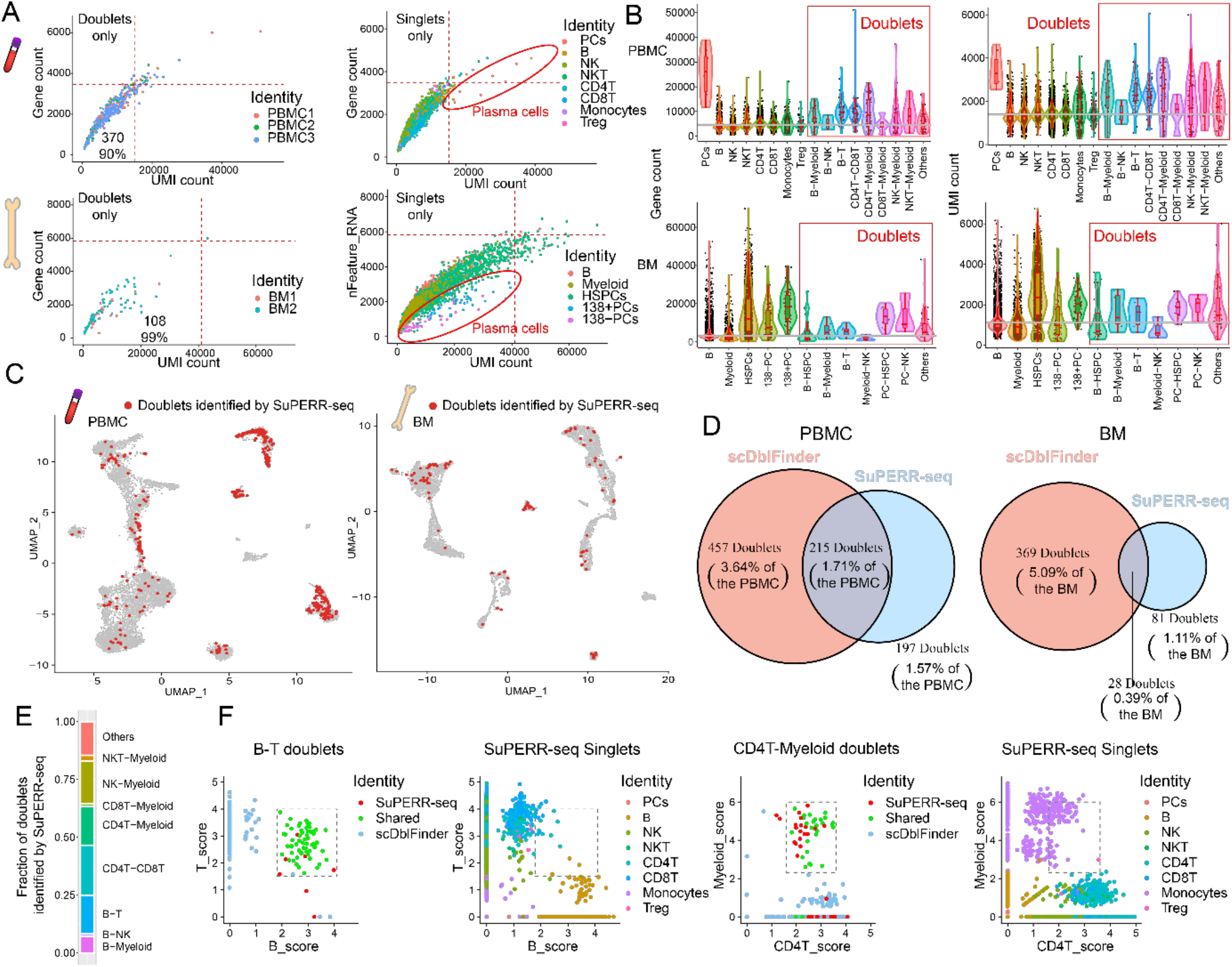
Cell-doublet identification by SuPERR-seq using both surface markers and gene expression data matrices. (A) Distribution (gene count x total UMI) of cell doublets (left) and singlets (right) detected by the SuPERR-seq approach. The red dashed lines show the threshold used by some conventional approaches to exclude cells that express higher than mean+4SD of gene count and total UMI. Only the cells above the dashed lines would have been excluded from the downstream analysis in conventional approaches (i.e., plasma cells in PBMCs, highlighted in the red circle, would have been incorrectly excluded from downstream analysis). (B) The number of unique genes (left panels) and the number of total UMIs (right panels) expressed by singlets and doublets in PBMC (top panels) and BM (bottom panels). The grey line shows the mean expression level across all clusters. (C) Cell doublets identified by the SuPERR-seq workflow and projected on a UMAP, showing the cell doublets are spread across multiple clusters. (D) Venn diagram comparing the cell doublets identified by the SuPERR-seq workflow and the ScDblFinder pipelines. (E) Proportion of heterotypic doublets identified and classified by SuPERR-seq in PBMC. (F) Expression level of gene signatures (see Methods) of heterotypic doublets defined by SuPERR-seq and scDblFinder to confirm their cell identities. Red points represent SuPERR-seq-defined doublets. Green points are the cell doublets identified by both SuPERR-seq and scDblFinder. Blue points represent scDblFinder-defined doublets, which were identified as singlets by SuPERR-seq. The immune cell types were annotated by the SuPERR-seq workflow.

We further compared our approach with current computational methods specifically designed to remove cell doublets in scRNA-seq datasets. First, we tested the standard workflow of scDblFinder (Germain et al. 2021) and compared it with the cell doublets defined by the SuPERR-seq method (Fig. 6D). To validate the cell doublets identified by both SuPERR-seq and scDblFinder doublets, we used a set of well-defined gene markers to calculate a cell-type score (see Methods) and then projected these scores for each cell barcode identified as doublets (Figs. 6E and 6F). We showed that SuPERR-seq and scDblFinder had good agreement on doublets that expressed high scores for multiple major lineages. Although scDblFinder outputted a larger number of cell doublets compared to the SuPERR-seq workflow, the scDblFinder-specific doublets did not show heterotypic patterns when compared against the singlets defined by both methods (Fig. S12), indicating potential false positives in the scDblFinder workflow. Most importantly, SuPERR-seq identified hundreds of validated cell doublets (i.e., cell barcodes that co-express markers, both mRNA and cell-surface protein, of more than one cell lineage) that were missed by the scDblFinder workflow (Fig. 6D).

### Benchmarking of SuPERR-seq against commonly-used methods reveal reduced cell-type misclassification, and superior ability to resolve and classify new cell types

We first compared our SuPERR-seq workflow with the commonly-used Seurat package v3 (Stuart et al. 2019) using default parameters. In the PBMC sample, Seurat v3 identified 24 clusters compared to 38 clusters identified by the SuPERR-seq workflow (Fig. S13A). In the BM sample, Seurat v3 identified 22 clusters compared to 34 clusters identified by SuPERR-seq (Fig. S14A). When we directly compared the cluster assignment of each cell barcode (Figs. S13A and S14A), multiple Seurat v3 clusters were further subdivided by the SuPERR-seq workflow into several biologically-meaningful clusters (Figs. 5 and S4-S11). The ability of the SuPERR-seq workflow to identify greater and novel subsets of immune cells can be explained by the advantages of pre-gating major immune cell lineages based on cell-surface markers and Ig-specific transcripts before exploring the GEX matrix. As shown in Fig. 4, pre-gating major lineages reveals the gene variation that occurs only *within* the pre-defined lineage instead of gene variation observed *across all* major lineages (Fig. 4D).

Importantly, our benchmarking analysis reveals a significant number of cell-type misclassifications (i.e., a single cluster containing cells from different major lineages) generated by the default Seurat v3 workflow. For example, Seurat v3 generated three clusters (5, 9, and 15) containing a mixture of cell lineages, including CD8+ T cells and NK cells (Fig. S13). In contrast, SuPERR-seq correctly clustered and classified these cells separately. Similarly, Seurat v3 cluster 3 mixed CD4+ and CD8+ T cells, while SuPERR-seq correctly identified and separated these different cell types (Fig. S13). Notably, such cell-type misclassification artifacts are not rare and commonly occur when simultaneously clustering all cells in high-dimensional space (Orlova, Herzenberg, and Walther 2018; Altman and Krzywinski 2018) using the GEX data matrix as performed by most, if not all, scRNA-seq analysis workflow.

One might attribute the improved performance of the SuPERR-seq workflow to the fact that it utilizes additional input data from other omics (i.e., ADT and V(D)J). Therefore, we further compared the SuPERR-seq workflow with two recently-developed pipelines that also integrate the information from both GEX and ADT data for clustering cells. We compared the SuPERR-seq workflow to the weighted nearest neighbor (WNN) approach as implemented in Seurat v4 (Hao et al. 2021) and the similarity network fusion (SNF) as implemented in CiteFuse (Kim et al. 2020) (Figs. S13B-C and S14B-C). To better quantify the performance of each data analysis workflow and to reveal the extent of unwanted cell-type misclassifications from each approach, we developed a new scoring system named Cell Fidelity Statistics (CFS) score (Babcock et al. 2021). Briefly, we consider the biaxial feature plots of cell-surface markers (ADT) and V(D)J-derived Ig-transcript counts (Figs. S1 and S2) to represent a “gold standard” as this iterative nature of SuPERR-seq prohibits cells from inappropriately co-clustering with a separate lineage. This approach is borrowed from the well-established biaxial gating strategy of flow cytometry analyses(Mahnke, Chattopadhyay, and Roederer 2010). We then compare the cell-type identity assigned to each cell barcode to those from a different workflow and consider the proportion of cells that change identity, generating a cell fidelity metric. The proportion of misclassified cells is reported as a “1-CFS score,” which provides a statistical measure of uncertainty in the cell-type assignment steps of the compared workflow. For example, the PBMC and BM clusters generated by Seurat v3 showed a 1 - CFS score of 0.0940 and 0.0531, respectively, indicating that 9.40% of PBMCs and 5.31% of BM cells were misclassified by the Seurat v3 (Fig. 7A-B). Notably, the “1-CFS scores” for the WNN (Seurat v4) and SNF (CiteFuse) approaches were lower than Seurat v3, indicating better agreement between Seurat v4, CiteFuse and SuPERR-seq and further highlighting the benefits of integrating additional omics for single-cell analysis (Fig. 7A-B). However, it is important to note that, similar to Seurat v3, WNN and SNF approaches still generated cell-type misclassifications (Fig. 7).

**Figure 7.**
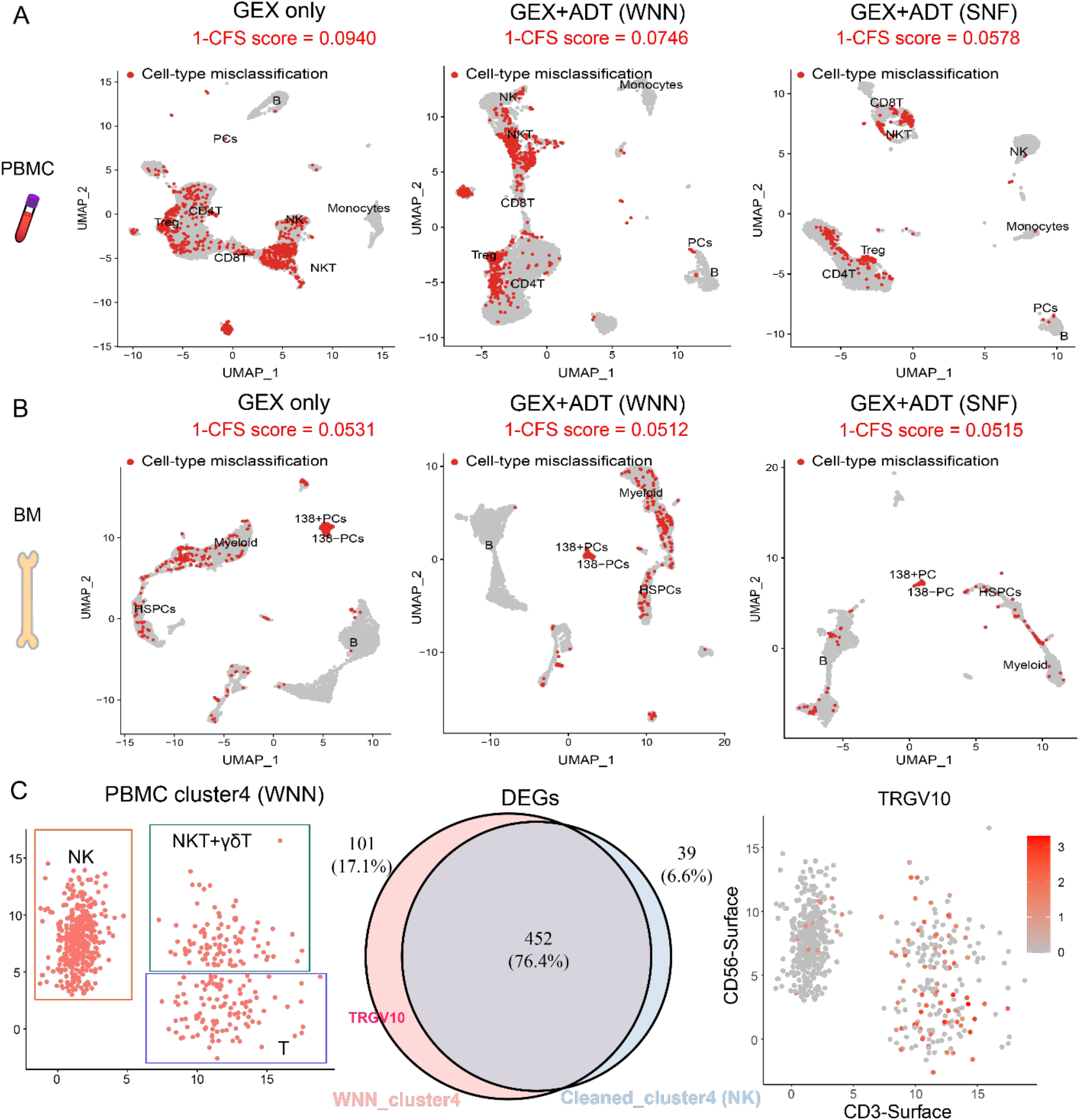
SuPERR-seq identifies significant cell-type misclassification from other approaches. (A) Red points represent the peripheral blood mononuclear cells (PBMC) that were misclassified by either the conventional approach using GEX data only (i.e., Seurat v3), or by more recent approaches using both GEX and ADT data, such as the WNN in Seurat v4, and the SNF in CiteFuse. The Cell Fidelity Statistic (CFS, see Methods) reports the fraction of correctly classified cells, the inverse of which is the fraction of misclassified cells (9.40% by Seurat v3, 7.46% by Seurat v4, 5.78% by CiteFuse). (B) Red points represent the bone marrow (BM) cells that were misclassified by Seurat v3 (5.31%), WNN/Seurat v4 (5.12%), and SNF/CiteFuse (5.15%) as determined by CFS. CFS scores show a progressive improvement in cell-type classification from Seurat v3 (GEX only) to Seurat v4 and CiteFuse, revealing higher agreement between CiteFuse and gold-standard biaxial gating of cell lineages. (C) The PBMC cluster 4 generated by the WNN method (Seurat v4) contains misclassified cells (i.e., a mixture of NK, NKT, and T cells) and was further explored using the cell-surface (ADT) markers CD56 and CD3 (left panel). The differentially expressed gene (DEG) analysis for cluster 4 (pink circle) compared to “cleaned” NK cells (Venn diagram) shows the *TRGV10* gene as a top hit. However, the *TRGV10* gene is mostly expressed in CD3^+^ gamma-delta T cells and absent in NK cells (right panel).

To further compare the performance of SuPERR-seq against WNN, we intentionally increased the clustering resolution in Seurat v4 from 0.5 to 3 to generate a total number of clusters that surpass the clusters generated by the SuPERR-seq workflow. For example, the default Seurat v4 pipeline at a cluster resolution 0.5 generated 34 and 29 clusters versus 38 and 34 clusters generated by SuPERR-seq in the PBMC and BM, respectively. In contrast, at a higher cluster resolution of 3, WNN generated 40 and 36 clusters in the PBMC and BM, respectively. Our rationale was that by generating additional clusters from the WNN approach we could observe improved cluster agreement with SuPERR-seq. However, even at higher cluster resolution and larger number of clusters, WNN was not able to identify a plasma cell cluster in the PMBC, and could not distinguish plasma cell subclusters in the BM (Figs. 7A-B, S13B and S14B). Similarly, the SNF approach (CiteFuse workflow) was not able to identify plasma cells in the PBMC, nor plasma-cell heterogeneity in the BM even after manually increasing the k (which is similar to increasing cluster resolution) for their spectral clustering approach. These results further support the SuPERR-seq workflow and its ability to generate biologically-meaningful subclusters of cells while preventing cell-type misclassifications.

To determine whether cell-type misclassifications generated by the other approaches could confound interpretation of downstream differential gene expression nalysis, we explored the PBMC cluster 4 generated by the WNN approach (Seurat v4) (Fig. 7C). Because DGE analysis is a commonly-used method to interpret biological differences between clusters or cell types, we hypothesized that running DGE analysis on a WNN (Seurat v4) cluster containing cell-type misclassification (as defined by SuPERR-seq) could generate misleading results even at low contamination (i.e., cell-type mixing) numbers. For example, the WNN-derived PBMC cluster 4 contains mainly NK cells, but it is also contaminated/mixed with NKT and T cells. Indeed, by analyzing the DGE list of the cluster 4 before removing the contaminating (NKT and T) cells, the *TRGV10* gene appeared as highly expressed for this cell population (Fig. 7C). However, *TRGV10* is expressed on gamma-delta T cells (Aliseychik et al. 2020), not on NK cells. These results indicate that the widespread cell-type misclassification that is often observed in conventional data-analysis workflows can confound data interpretation.

Finally, when comparing the new SuPERR-seq-identified plasma cells with the other workflows, we observed major differences. While SuPERR-seq readily and accurately identified a cluster of eight plasma cells in PBMCs (Fig. 2A), the other pipelines (Seurat v3, WNN/Seurat v4, and SNF/CiteFuse) failed to identify these rare cells (Fig. S13). Unlike PBMC, BM cells contain significantly more plasma cells. In this case, SuPERR-seq identified four biologically distinct plasma cell clusters, while the other workflows identified only one cluster (Figs. 5 and S14). Of note, the plasma cell cluster generated by the Seurat v3 workflow contained a mixture of cell types (i.e., cell-type misclassification), including B cells and HSPCs (Fig. S14A). In addition, all the workflows we tested have misclassified a fraction of the SuPERR-seq-identified plasma cells, as they appeared spread across multiple clusters (Fig. S14).

To determine whether further optimizations in the analysis workflow would allow the other approaches (Seurat v3, WNN and SNF) to identify the four SuPERR-seq plasma-cell clusters in the BM, we performed a second iteration of clustering (i.e., recursive approach/re-clustering) on the original BM plasma cell cluster identified by both Seurat (v3 and v4) and CiteFuse. After the second iteration of clustering, Seurat (v3 and v4) and CiteFuse were able to identify more BM plasma cell clusters (Fig. S15). However, the plasma cells clusters remained as a mixture of contaminating B cells and HSPCs (Fig. S15). For the fairness of comparison, the parameters used for running the Seurat v3, Seurat v4, CiteFuse and SuPERR-seq workflows were set as the default parameters described in Methods (unless noted otherwise). And the re-clustering approach on the original Seurat v3, Seurat v4 and CiteFuse clusters was limited to two iterations.

Taken together, the results from the benchmarking analysis show that the SuPERR-seq workflow not only identifies more and novel clusters but also improves the clustering purity and the accuracy of biological interpretation.

## Discussion

A core component of a single-cell RNA-seq analysis is the identification and classification of cell types. The most popular and widely-used approaches rely on principal component analysis (PCA) derived from a set of highly variable genes (HVGs), followed by nearest-neighbors graph-based clustering. The resulting clusters are then assigned to a cell type and biological function via manual annotation, using the mRNA transcript counts as a guide. In existing approaches, the selection of HVGs for downstream clustering analysis is often dominated by lineage-specific markers. These markers often capture variance that exists *between* cell lineages (e.g., the variance that distinguishes T cells from B cells) but often misses variance that exists *within* each cell lineage (e.g., the variance that distinguishes the various T-cell subsets, including effector memory from central memory, stem memory, naïve, etc.) (Fig. 4). Thus, an improved method to select HVGs *within* each cell lineage is needed to discover new biologically-meaningful cell subtypes with high accuracy.

Moreover, selecting HVGs with high variance across all lineages (instead of *within* lineages) as the very first step in data analysis (as in many existing approaches) carries the risk of inappropriately grouping cells with similar gene expression but arising from separate cell lineages, resulting in cell-type misclassification (Fig. 7). Cell-type misclassification often occurs in conventional scRNA-seq data analyses. It can be caused by a mathematical phenomenon known as the curse of dimensionality (Altman and Krzywinski 2018; Trevor Hastie 2009; Orlova, Herzenberg, and Walther 2018), inadvertently misguiding biological interpretations (Fig. 7, 1-CFS scores). The integration of additional omics, including cell-surface markers used in WNN (Hao et al. 2021) and SNF (Kim et al. 2020) approaches, can improve cell-type classification, however, it doesn’t eliminate cell-type misclassification. Another contributing factor to cell-type misclassification is the presence of cell doublets in the final dataset. Despite recent advances in microfluidics that precisely generate droplets containing single cells, a significant fraction of the droplets can still contain more than one cell (Allon et al. 2015; Cao et al. 2017). Current approaches to remove cell doublets rely on overly high mRNA transcript counts (i.e., outliers) (Ocasio et al. 2019) and gene marker co-expression. However, as we show here, many cell doublets contain average mRNA counts and can mistakenly be carried over to downstream analysis (Fig. 6). More recent and sophisticated computational algorithms to identify cell doublets such as scDblFinder (Germain et al. 2021) can still ignore true doublets and captures false positives (Figs. 6 and S12).

To address these pitfalls of existing methods, we presented here the semi-supervised SuPERR-seq workflow. In SuPERR-seq, we simultaneously apply information gained from triple-omics sequencing, namely gene expression (GEX), cell-surface proteins measured by antibody-derived tags (ADT), and immunoglobulin transcript counts from the V(D)J repertoire matrix. As a quality control measure and to prevent downstream cell-type misclassification, the first step of SuPERR-seq is to perform a “manual gating” similar to the standard flow cytometry analysis. ADT and V(D)J matrices can reliably classify cells into clearly distinct major lineages using well-established canonical markers. This manual gating step ensures that any downstream subclusters are composed of a single lineage, allowing for a more accurate cell-type classification and data interpretation. The second step of the SuPERR-seq is to examine each lineage independently, both from one another and from the first gating steps. Rather than selecting genes with high variance across all cell lineages, we select HVGs only from within each manually-gated lineage. These HVGs inform a PCA and subsequent nearest-neighbors graph-based clustering. Our PBMC and BM samples analysis show that the SuPERR-seq approach can reveal additional cell types (and likely cell states) compared to other existing approaches. We reason that this is mainly due to the ability of SuPERR-seq to select only the relevant HVGs for each well-defined lineage and ignore extra sources of variance that may not be informative within each cell lineage.

To ensure the generalizability of our biaxial gating approach so that SuPERR-seq can be broadly applicable regardless of the type of sample/tissue, we included in our ADT panel well-established cell-surface markers that have been previously validated using the gold-standard flow cytometry approach (Fig. S3). These cell-surface markers can readily (and accurately) identify major cell lineages, including immune lineages (i.e., CD45+), non-immune lineages (i.e., CD45-), epithelial lineages (i.e., EPCAM+), and other lineages regardless of the origin of the sample. Of note, many lineage-specific cell-surface markers have been validated and published in open-source journals, including the collection of cell-type classifications available as Optimized Multicolor Immunophenotyping Panels (OMIPs) (Mahnke, Chattopadhyay, and Roederer 2010). These well-characterized cell-surface markers can be included as part of the ADT panels used in any experimental design to identify major immune and non-immune lineages. Thus, new datasets with different combinations of antibody (ADT) panels can still follow a similar SuPERR-seq gating strategy as long as the panel includes known lineage markers. Furthermore, we expect the SuPERR-seq approach to improve the ability to identify novel cell subsets and cellular states (within each major lineages) for which cell-surface markers have not yet been characterized. For example, a novel subset (cluster) of B cells can be discovered within the major B-cell lineage (i.e., within cells expressing surface CD19). In the (unlikely) event that the novel B-cell subset doesn’t express CD19, the novel subset should still form a separate cluster within another pre-defined gate based on its cell-surface marker expression. In this scenario, the additional omics (GEX and VDJ) in our workflow would reveal the B-cell identity of the novel B-cell subset.

Future studies to optimize the initial biaxial manual gating step might include developing an automated semi-supervised clustering algorithm to readily identify major cell lineages using the ADT and V(D)J data matrices. Automated algorithms that still rely on biaxial projections, such as the Exhaustive Projection Pursuit (EPP) approaches (Friedman and Tukey 1974) might provide more accurate results compared to fully unsupervised methods in high-dimensional space, which might suffer from the curse of dimensionality (Altman and Krzywinski 2018; Trevor Hastie 2009; Orlova, Herzenberg, and Walther 2018). In fact, we recently implemented an automated EPP approach to identify cell subsets using cell-surface markers in high-dimensional flow and mass cytometry datasets (Meehan et al. 2019). Although this new Subset Identification and Characterization (SIC) pipeline was specifically developed for flow and mass cytometry datasets (Meehan et al. 2019), future studies should aim to optimize such pipelines to process multi-omics single-cell ADT and V(D)J matrices as an alternative automated step in the SuPERR-seq workflow.

We recognize the difficulty and expense of generating triple-omics data for every sample, as well as the limitations of applying the SuPERR-seq approach to older data sets for which ADT and V(D)J matrices are unavailable. Also, some samples might not include B and/or T cells, which means there will not be V(D)J matrices. Thus, to broaden the application of the SuPERR-seq principles to datasets without triple-omics available, we propose a gene-based (GEX) recursive analysis approach (i.e., re-clustering), which carries some of the same benefits of SuPERR-seq. When we applied this recursive approach to the same BM dataset explored by the SuPERR-seq we identified improvements, but also limitations (Fig. S15). Focusing on the plasma cells, the recursive approach applied to other existing workflows, which only identified one cluster of plasma cells in the first clustering iteration, now generated more subclusters of plasma cells. However, the re-clustering approach was not able to accurately identify biologically-meaningful plasma cell clusters and, most importantly, it was not able to isolate the plasma cells from the cell contaminants. Thus, the resulting subclusters of plasma cells represented a mixture of plasma cells, B cells, and HSPCs (Fig. S15). In contrast, SuPERR-seq readily and accurately identified four subsets of plasma cells that were confirmed based on the three omics (GEX, ADT, VDJ), including the high antibody transcript counts (∼2.5 log_10_-fold higher antibody transcripts than B cells). Moreover, the plasma cell subsets identified by SuPERR-seq are of great biological importance as revealed by the Reactome pathway analysis (Fig. 5). Therefore, integrating three omics in the SuPERR-seq workflow provides unique information that recursive clustering strategies for defining HVGs cannot achieve. Our conclusions here agree with the latest implementation of Seurat v4 (Hao et al. 2021) and CiteFuse (Kim et al. 2020), which also integrate ADT measurements in their analysis pipeline, further supporting the SuPERR-seq approach of multi-omics integration for defining cell clusters.

Taken together, we developed a comprehensive multi-omics single-cell data integration and analysis workflow that mitigates or resolves pitfalls in existing approaches and allows for the discovery of novel and biologically-meaningful cell subsets in the human immune system.

## Methods

### Subjects and specimen collection

Peripheral blood was collected from healthy adult donors through Emory University’s Children’s Clinical and Translational Discovery Core (CCTDC). Bone marrow samples were collected from healthy adult donors through Emory University Hospitals under IRB00066294 or obtained from AllCells (Alameda, CA).

### Cell preparation

Peripheral blood mononuclear cells (PBMCs) were isolated from peripheral venous blood using EasySep™ Direct Human PBMC Isolation Kit (Cat # 19654). B cells were enriched from hip bone marrow (BM) using EasySep™ Direct Human PBMC Isolation Kit (Cat # 19654) followed by EasySep Human B-Cell Enrichment Kit II without CD43 Depletion (Cat # 17963) following manufacture’s protocol. Up to 1X10^6^ cells per donor were incubated with Fc block (Miltenyi) on ice for 10 min followed by staining with a mix of 31 oligo-conjugated (barcoded) antibodies (see Table S1 for full list) for 30 min on ice, in custom-made RPMI-1640 media deficient in biotin, L-glutamine, phenol red, riboflavin, and sodium bicarbonate, and containing 3% newborn calf serum (def-RPMI-1640), followed by two washes in def-RPMI-1640/0.04% BSA. Cells were resuspended at a concentration of 1200-1500 cells/uL in custom RPMI-1640/0.04% BSA and passed through a 20-um nylon filter before loading onto a Chromium Controller (10X Genomics, Pleasanton, CA).

### Single-cell encapsulation, library generation, and sequencing

Cells were loaded to target encapsulation of 6,000 cells. Gene expression (GEX), Antibody-derived tag (ADT), and V(D)J libraries were generated using the Chromium Single Cell 5′ Library & Gel Bead Kit v1 with feature barcoding (10X Genomics, Pleasanton, CA) following the manufacturer’s instructions. Gene expression libraries were pooled and sequenced on a NovaSeq 6000 platform (Illumina, San Diego, CA). ADT and V(D)J libraries were sequenced separately on a Next-seq platform (Illumina, San Diego, CA). The sequencing depths are shown in Table S2.

### Multi-omics single-cell data preprocessing

10X Genomics CellRanger v3.1.0 was used to perform barcode processing and single-cell 5′ unique molecular identifier (UMI) counting. Reads from GEX and ADT libraries were processed simultaneously using “cellranger count,” while reads from V(D)J libraries were aligned by running “cellranger vdj.”

### Gene expression (GEX)

The scRNA-seq expression datasets were integrated into two final matrices, one for PBMC and another for BM. To remove potential batch effects, we used the canonical correlation analysis (CCA) to integrate the three individual PBMC samples into one final PBMC dataset and combine the two individual BM samples into one final BM dataset. Before integration, we performed library-size scaling and log-transformation for each sample individually. Next, we identified the top 2,000 highly variable genes for each sample using regularized negative binomial regression (Hafemeister and Satija 2019), followed by CCA to find anchors and integrate cells from individual samples into one integrated dataset. Finally, we performed per-gene z-score normalization for each integrated dataset. We removed from the downstream analysis the genes expressed in less than three cells, the cells with less than 200 genes, and the cells with more than 20% of mitochondrial gene counts. GEX raw count matrices are available through NCBI GEO, accession number GSE181543.

### Antibody-derived tags (ADT)

The cell-surface protein expression (ADT) data were normalized using the DSB normalization (Mulè, Martins, and Tsang 2021). Cell barcodes in the CellRanger unfiltered matrix containing less than 40 genes and less than 100 total UMIs were considered background. The barcodes from CellRanger “filtered_feature_bc_matrix” were considered true cells. By subtracting the mean value of the background population and then regressing out the cell-cell technical variation (Mulè, Martins, and Tsang 2021), the ADT values representing background expression levels were centered around zero. ADT raw count matrices are available through NCBI GEO, accession number GSE181543.

### Antibody repertoire (VDJ)

We used the CellRanger-generated “all_contig_annotations.csv” (unfiltered) and the “filtered_contig_annotations.csv” files to generate new V(D)J features merged with the corresponding ADT matrix. Briefly, we summed the total UMIs of each cell barcode in the “all_contig_annotations.csv” V(D)J file to generate a feature called “Total Ig Transcripts” that represent the total UMIs of immunoglobulin heavy and light chains. In addition, we generated a binary feature called “Productive VDJ” that identifies whether a cell barcode with a productive V(D)J information is contained in the CellRanger “filtered” annotation file and hence considered a true B cell or plasma cell. B-cell receptor repertoire V(D)J raw count matrices are available through NCBI GEO, accession number GSE181543.

### Manual gating

Manual gating was performed based on the two V(D)J features (i.e., Ig-specific transcript counts and productive VDJ sequences) and the 31 cell-surface protein features in the normalized ADT data. The major cell lineages (six for PBMCs and five for BM cells) were identified and manually gated using a customized strategy of biaxial plots implemented in a MATLAB script (https://github.com/Ghosn-Lab/SuPERR-seq.git). The markers and the gating hierarchy were determined by prior knowledge of variations of cell-surface protein expression across various major cell lineages (Figs. 2A and 3A). Cell doublets were manually “gated out” based on the co-expression of two or more major lineage markers (Figs. S1 and S2).

### Clustering on gene expression matrix

Each major cell lineage identified by the manual gating was next clustered based on the integrated GEX matrix (Stuart et al. 2019). First, HVGs were computed for cells in each major cell lineage using the variance-stabilizing transformation (VST) method (Hafemeister and Satija 2019), and expression data of each gene was transformed to zero mean and unit variance. We then performed PCA on the highly variable genes of each main lineage and selected the top 30 PCs for downstream analysis. Next, a K-nearest neighbor (KNN) graph was constructed in the low-dimensional PCA space based on the Euclidean distance between cells, with K=30. Jaccard further converted the KNN graph to a cell-cell similarity matrix, followed by Louvain community detection algorithm (Blondel et al. 2008) to define cell clusters in each major cell lineage.

The weighted nearest neighbor (WNN) approach as implemented in Seurat v4 and the similarity network fusion (SNF) as implemented in CiteFuse were separately applied to generate cell clusters cells based on an intergrated matrix containing both GEX and ADT data. For Seurat v4, we used the graph-based smart local moving (SLM) algorithm (Waltman and van Eck 2013), and set the clustering resolution to 3 to generate as many (or more) clusters as SuPERR-seq. For CiteFuse, cells from different samples were first integrated using CCA as described above and then the integrated dataset was used as input. We applied the spectral clustering algorithm as suggested by the authors and the K was set to 39 for PBMC and 35 for BM to align the number of cell clusters generated by SuPERR-seq.

### Differential gene expression analysis

We used the “FindAllMarkers” function of Seurat v3 to perform the differential gene expression analysis. Statistical significance was tested using the Wilcoxon Rank-Sum test, with p_val_adj<0.05 (p-value after Bonferroni correction). For the DGE analysis of the BM plasma cell clusters, we first removed the immunoglobulin-specific UMIs from the GEX matrix. Then, we log-normalized the resulting matrix to discover and explore non-immunoglobulin plasma cell genes.

### Pathway analysis

The DGE list for each BM plasma cell cluster was used as input data to the Reactome pathway database (Jassal et al. 2020) (https://reactome.org/PathwayBrowser/) to visualize active biological pathways and regulatory processes in each plasma cell cluster. The “firework” diagrams were cropped so that only the hit pathways were preserved.

### Comparison of cell doublets identified by scDblFinder and SuPERR-seq

The scDblFinder (Germain et al. 2021) workflow was run independently on the GEX matrix from the PBMC and BM samples, following the workflow described on github (https://github.com/plger/scDblFinder). First, we performed log-normalization, HVG selection and PCA analysis on the GEX matrix for each sample. Next, we ran the scDblFinder pipeline using the default parameters.

### Calculation of cell-type scores

Each cell-type specific score was calculated by summing the raw UMI counts from the gene list we generated based on prior knowledge and log-normalized the data. The following gene lists were used for each cell-type score. *B cells*: CD79A, MS4A1, CD19, VPREB3; *CD4T*: CD4, CD3D, CD3E, CD5, IL7R; *CD8T*: CD8A, CD8B, GZMK, CD3D, CD3E; *Myeloid*: LYZ, S100A8, S100A9, S100A12, CD68, CD14, CYBB; *NK*: GNLY, NKG7, GZMB, KLRD1, GZMA; *NKT*: CCL5, GNLY, NKG7, GZMH, KLRB1, CST7; *Treg*: FOXP3, CTLA4, IL2RA, IL32; *T cells*: CD3D, CD3E, CD4, CD8A, CD8B, CD5, IL7R, GZMK, GZMH; and *PC* (Plasma cell score): ITGB7, IRF4, CD9, PRDM1, XBP1, SDC1, VCAM1, CD38.

### Cell Fidelity Statistic

To generate the Cell Fidelity Statistic (CFS) score (Babcock et al. 2021), we cross-check the cell lineages identities generated by a biaxial gating approach (reference) versus cell cluster assignment (test). CFS relies on the tenet that if different workflows identify a single cell as belonging to multiple lineages, both cannot be correct and therefore some information has been lost. We apply CFS to measure the loss in cell classification fidelity caused by relying on clustering algorithms to discriminate cell lineages, as is the norm in conventional workflows. To generate a CFS score, we first produce clusters (Seurat v3, Seurat v4, CiteFuse). We then count the number of cells which are grouped into majority out-of-lineage clusters. The CFS score is expressed as a fraction of total cells which were grouped into the inappropriate lineage, or the cells which have a disagreement between cluster assignment and biaxial gate. CFS is reported as a fraction from 0-1, where 1 means that no cells were misclassified, and a score of 0.8 means that 20% of cells were misclassified.

## Acknowledgments

This study was supported in part by Georgia Clinical and Translational Science Alliance (CTSA) through the National Center for Advancing Translational Sciences of the National Institutes of Health under Award number NIH UL1TR002378 (E.E.B.G.); Pediatric Research Alliance, Center for Transplantation, and Immune-Mediated Disorders (E.E.B.G.); Lowance Center for Human Immunology (E.E.B.G.); and the National Science Foundation awards CCF1552784 and CCF2007029 (P.Q.). P.Q. is an ISAC Marylou Ingram Scholar and a Carol Ann and David D. Flanagan Faculty Fellow. We thank Sachin Kumar (Emory University) for providing the Python script used to generate the Circos plots and Matthew C. Woodruff (Emory University) for the R script used to generate the V(D)J input files for the Circos plots. Flow cytometry data were collected at the Emory’s Pediatrics/Winship Flow Cytometry Core (access supported in part by Children’s Healthcare of Atlanta). Single-cell libraries were sequenced at the Emory Integrated Genomics Core (EIGC), which is subsidized by the Emory University School of Medicine and is one of the Emory Integrated Core Facilities; the Parker H. Petit Institute for Bioengineering and Bioscience at the Georgia Institute of Technology; and the PerkinElmer Genomics Inc. We thank Dr. F. Eun-Hyung Lee (Emory University), Sang N. Le, and Mindy R. Hernández for kindly providing the bone marrow aspirates from healthy adult donors through Dr. Lee’s Emory IRB protocol, and Emory University’s Children’s Clinical and Translational Discovery Core (CCTDC) for providing peripheral blood samples from healthy donors.

## Author Contributions

E.E.B.G. and P.Q. conceived the idea and experimental design, interpreted the data, and wrote the final draft. C.M. and J.Y. generated, analyzed, and interpreted the multi-omics single-cell datasets and generated the figures. J.Y. performed the preprocessing workflows to obtain final count matrices for each single-cell dataset. A.K. received and processed the human samples, generated the single-cell libraries and flow cytometry datasets, and analyzed the data. B.R.B. provided input into data analysis and generated the CFS scores. All authors contributed to the writing of the manuscript and approved the final version.

---

**Table S1.**
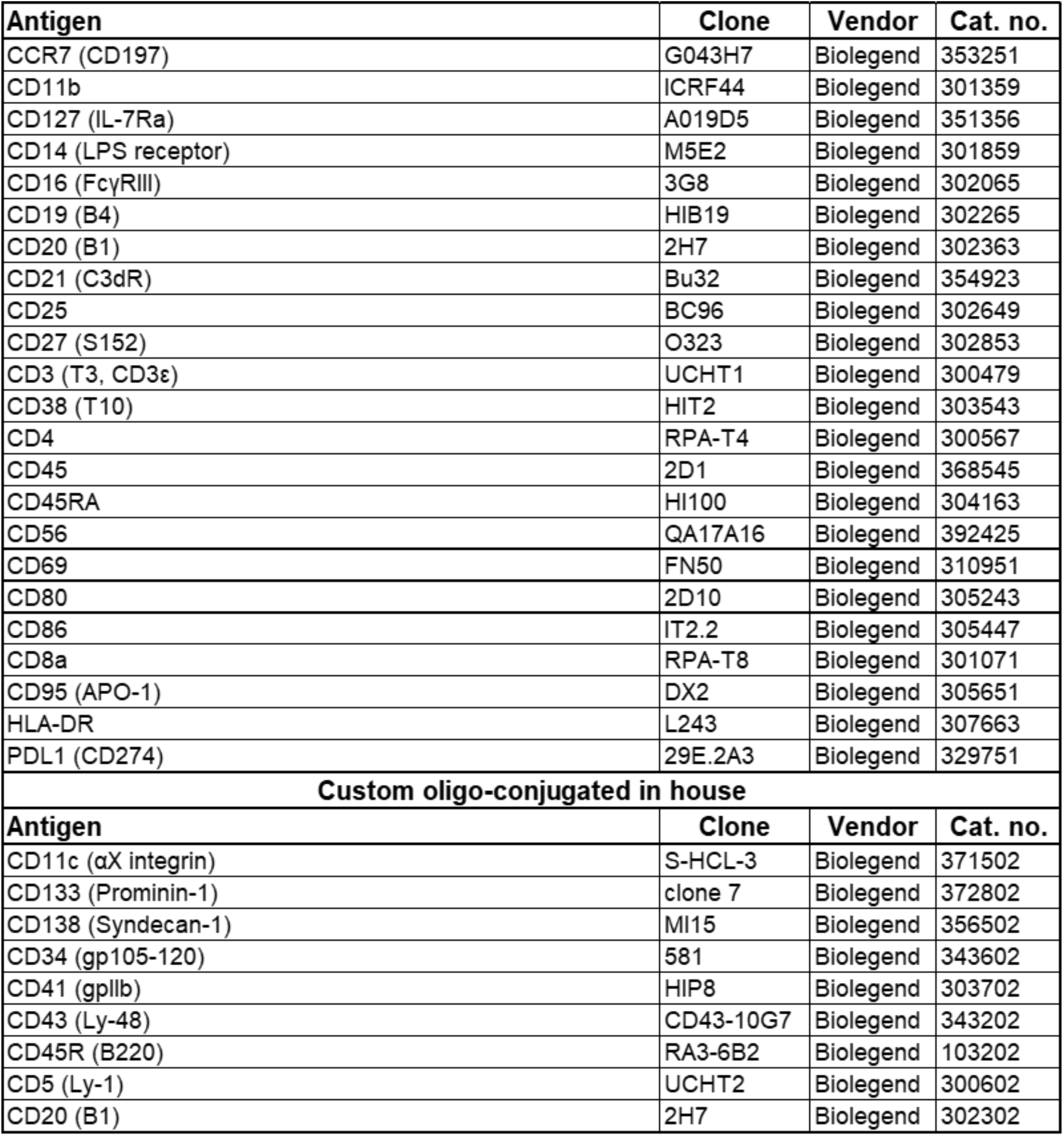
List of oligo-conjugated antibodies to determine antibody-derived tags (ADT) used in the multi-omics single-cell assays.

**Table S2.** Sequencing information. Mean reads per cell (R2 reads) and the total number of cells analyzed in each sample. PBMC: peripheral blood mononuclear cells; BM: bone marrow cells.

|  | Sample ID | Mean reads per cell | Cell count |
| --- | --- | --- | --- |
| Gene expression | PBMC1 | 138,274 | 4,034 |
|  | PBMC2 | 150,868 | 2,302 |
|  | PBMC3 | 45,805 | 6,423 |
|  | BM2 | 149,030 | 3,480 |
|  | BM3 | 149,622 | 3,946 |
| Surface protein expression | PBMC1 | 1,399 | 4,034 |
|  | PBMC2 | 2,273 | 2,302 |
|  | PBMC3 | 928 | 6,423 |
|  | BM2 | 2,762 | 3,480 |
|  | BM3 | 1,703 | 3,946 |
| V(D)J expression | PBMC1 | 126,518 | 273 |
|  | PBMC2 | 601,049 | 64 |
|  | PBMC3 | 56,584 | 582 |
|  | BM2 | 22,991 | 1,669 |
|  | BM3 | 17,418 | 2,553 |

**Figure S1.**
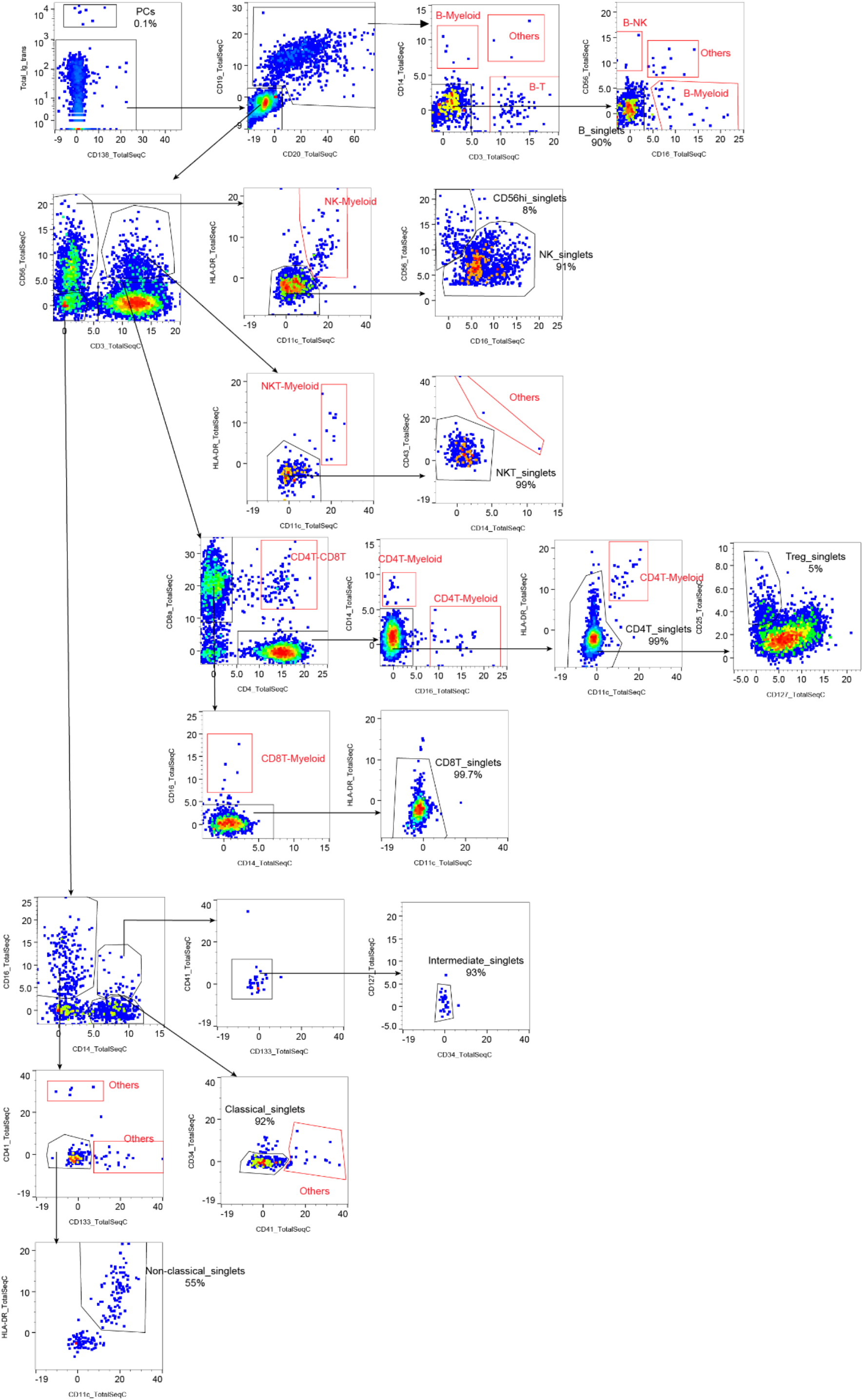
Biaxial feature plots (ADT data) to identify and remove cell doublets from PBMCs. Supplement for Figure 2. Cell doublets were gated (red boxes) based on co-expression of cell-surface markers (ADT) from two or more major lineages. The visualization was generated by SeqGeq^TM^.

**Figure S2.**
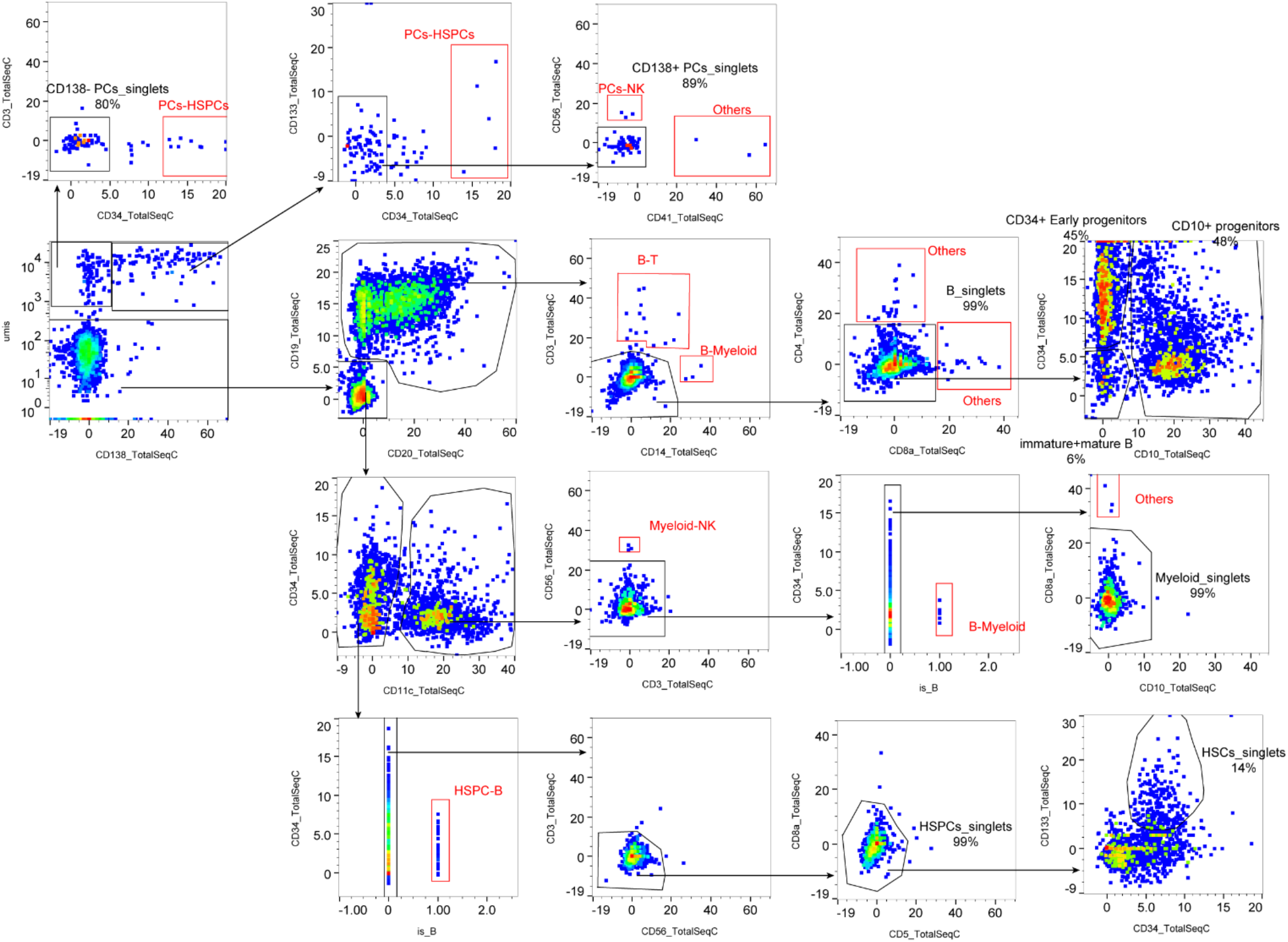
Biaxial feature plots (ADT data) to identify and remove cell doublets from BM cells. Supplement for Figure 3. Cell doublets were gated (red boxes) based on co-expression of cell-surface markers (ADT) from two or more major lineages. The visualization was generated by SeqGeq^TM^ (FlowJo, LLC).

**Figure S3.**
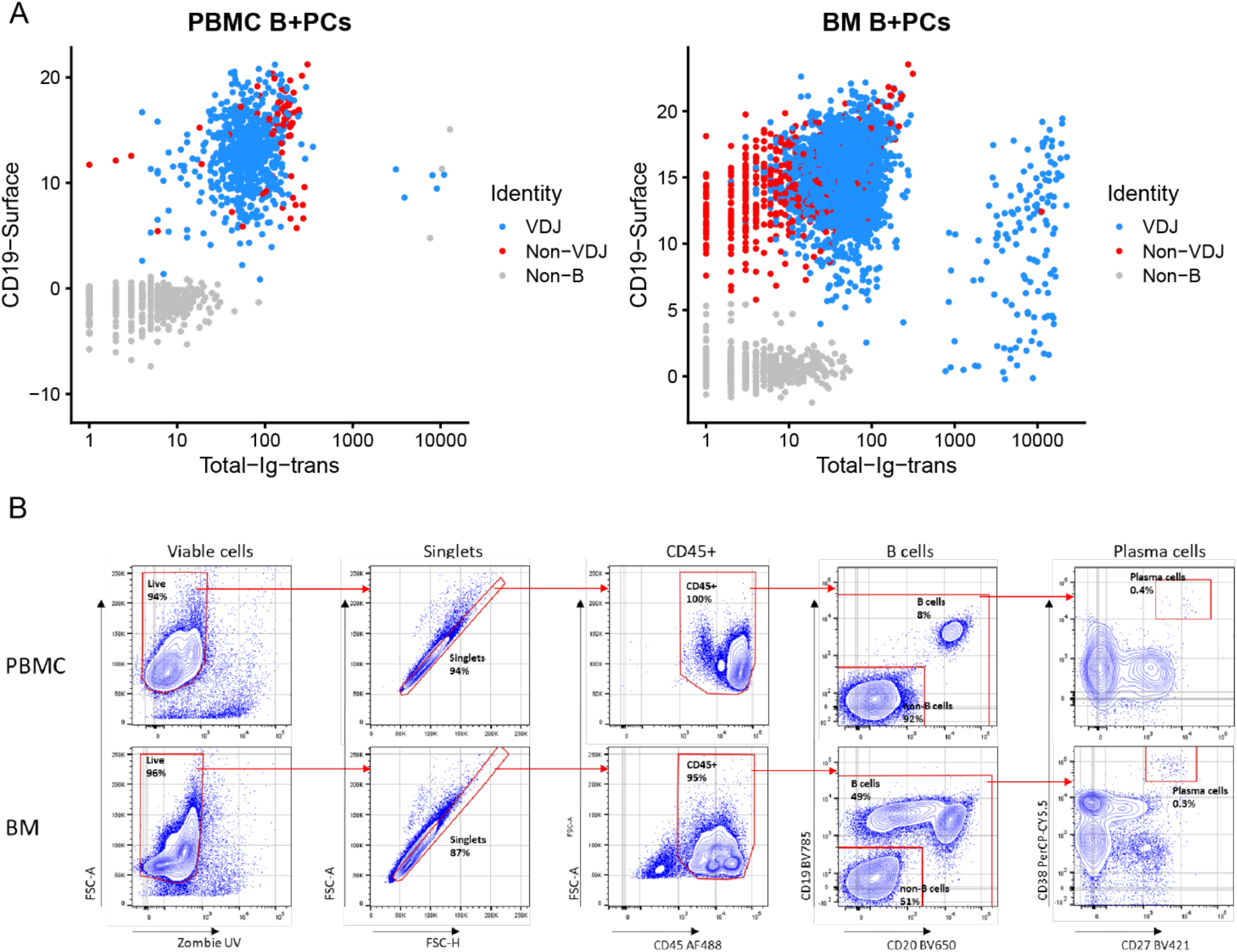
Validation of the SuPERR-seq approach of cell-type classification using ADT. (A) Bixal plot showing the expression level of CD19 ADT and Total Immunoglobulin transcripts (VDJ) of the scRNA-seq data. Grey points are non-B cells defined by SuPERR-seq workflow. Red points are B cells not included in the VDJ matrix. Blue point are B cells included in the VDJ matrix. (B) Gating strategy to identify major lineages by high-dimensional flow cytometry. An aliquot of each PBMC (top row) and BM (bottom row) sample was stained with a panel of fluorescently-labeled antibodies in parallel to the SuPERR-seq workflow. The gating strategy (red gates) shows the percentage of live, single B cells, and plasma cells.

**Figure S4.**
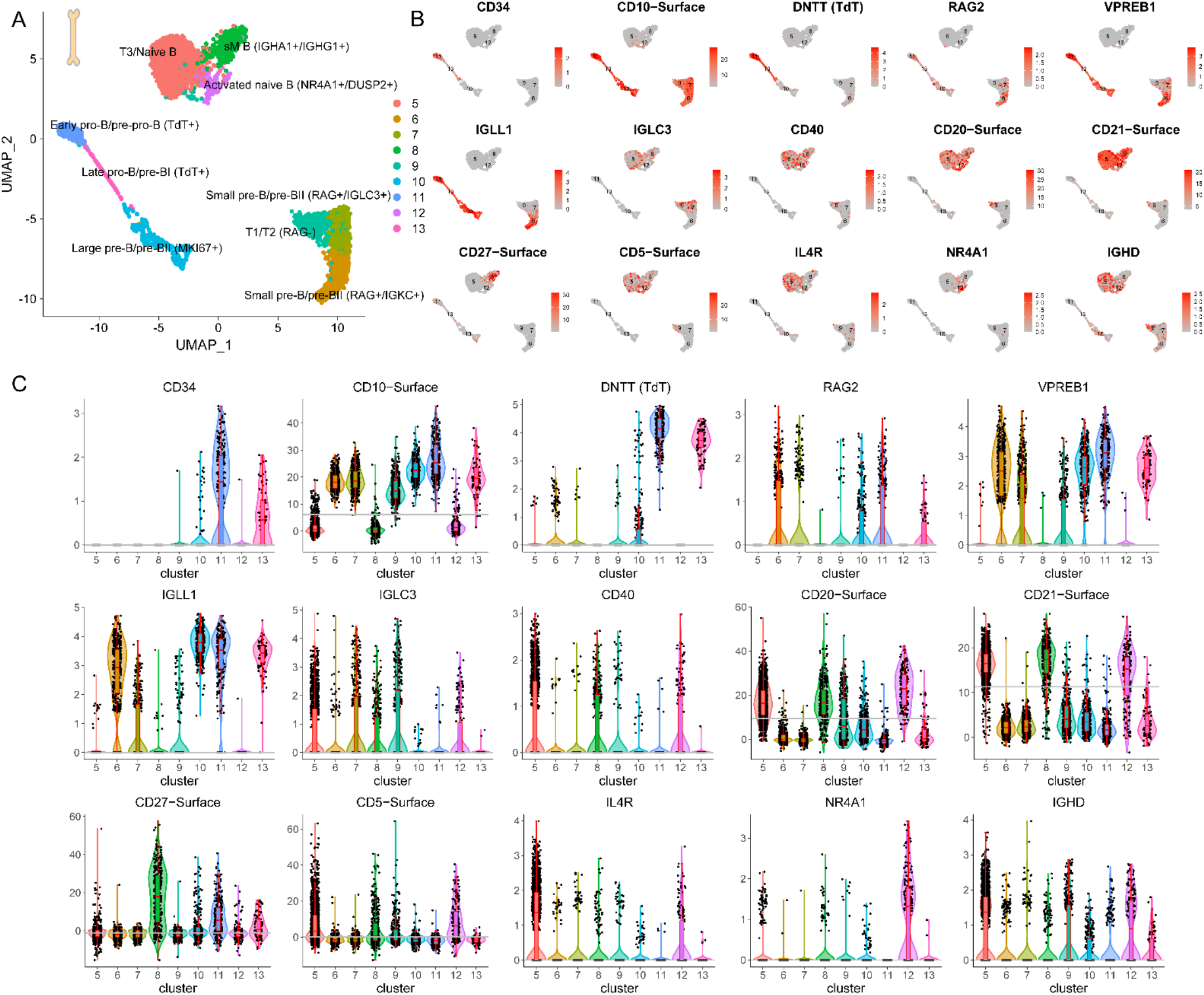
B-cell developmental pathway in human adult BM identified by the SuPERR-seq workflow. (A) Cell-type classification of bone marrow (BM) B cells based on three -omics: cell-surface protein (ADT), immunoglobulin heavy and light chain repertoire (VDJ), and transcriptomics/gene expression (GEX). (B) The expression level of cell-surface proteins (ADT) and genes (GEX) used to define the B-cell developmental stages. (C) Violin plots illustrate the differences in expression levels (ADT and GEX) between clusters. The grey line shows the mean expression level across all clusters.

**Figure S5.**
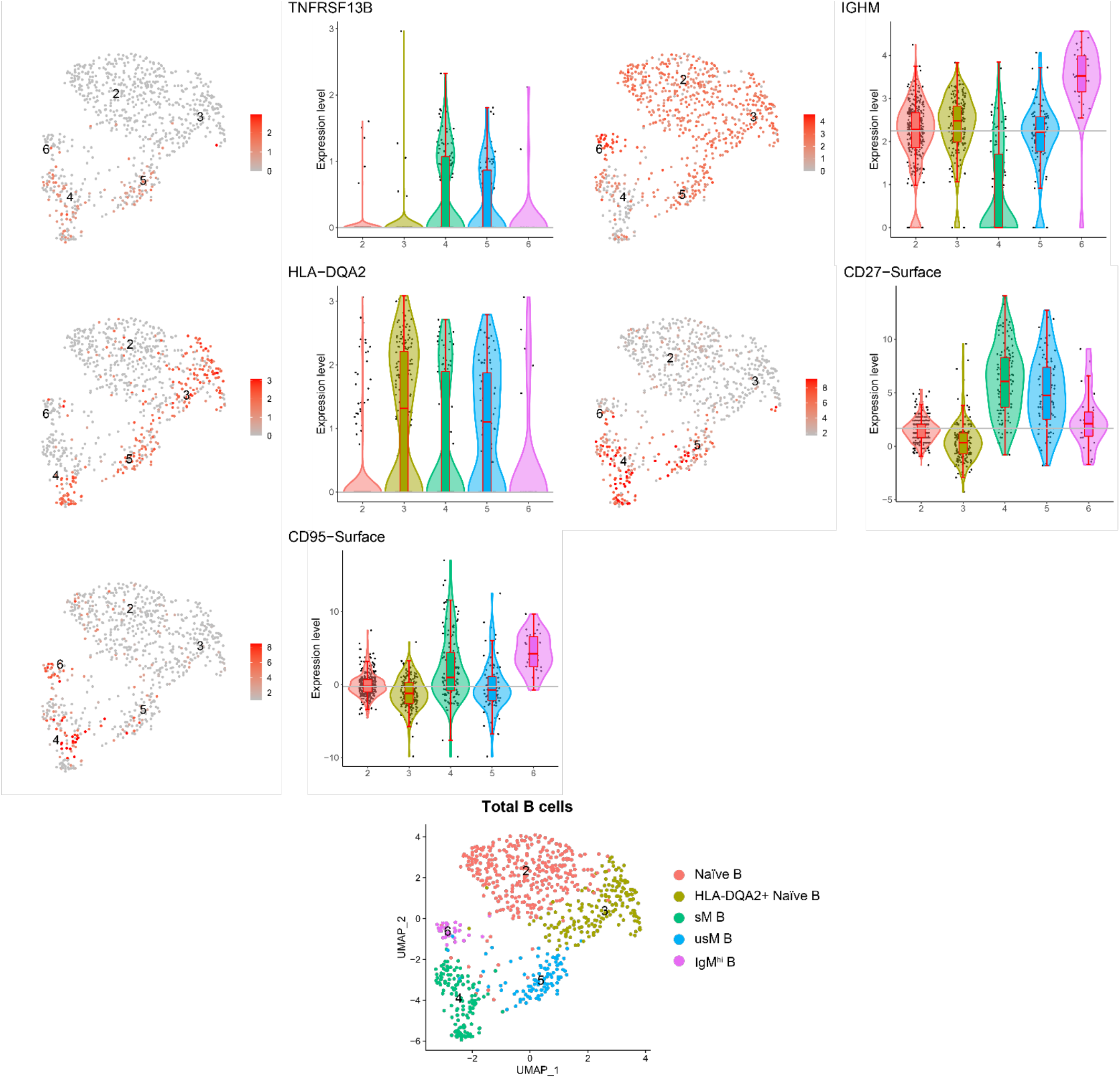
Blood B cells identified by the SuPERR-seq workflow. UMAP and violin plots show cell-surface markers (ADT) and gene expression (GEX) levels used to identify and classify five subsets of B cells in peripheral blood mononuclear cells (PBMC). The grey line shows the mean expression level across all clusters.

**Figure S6.**
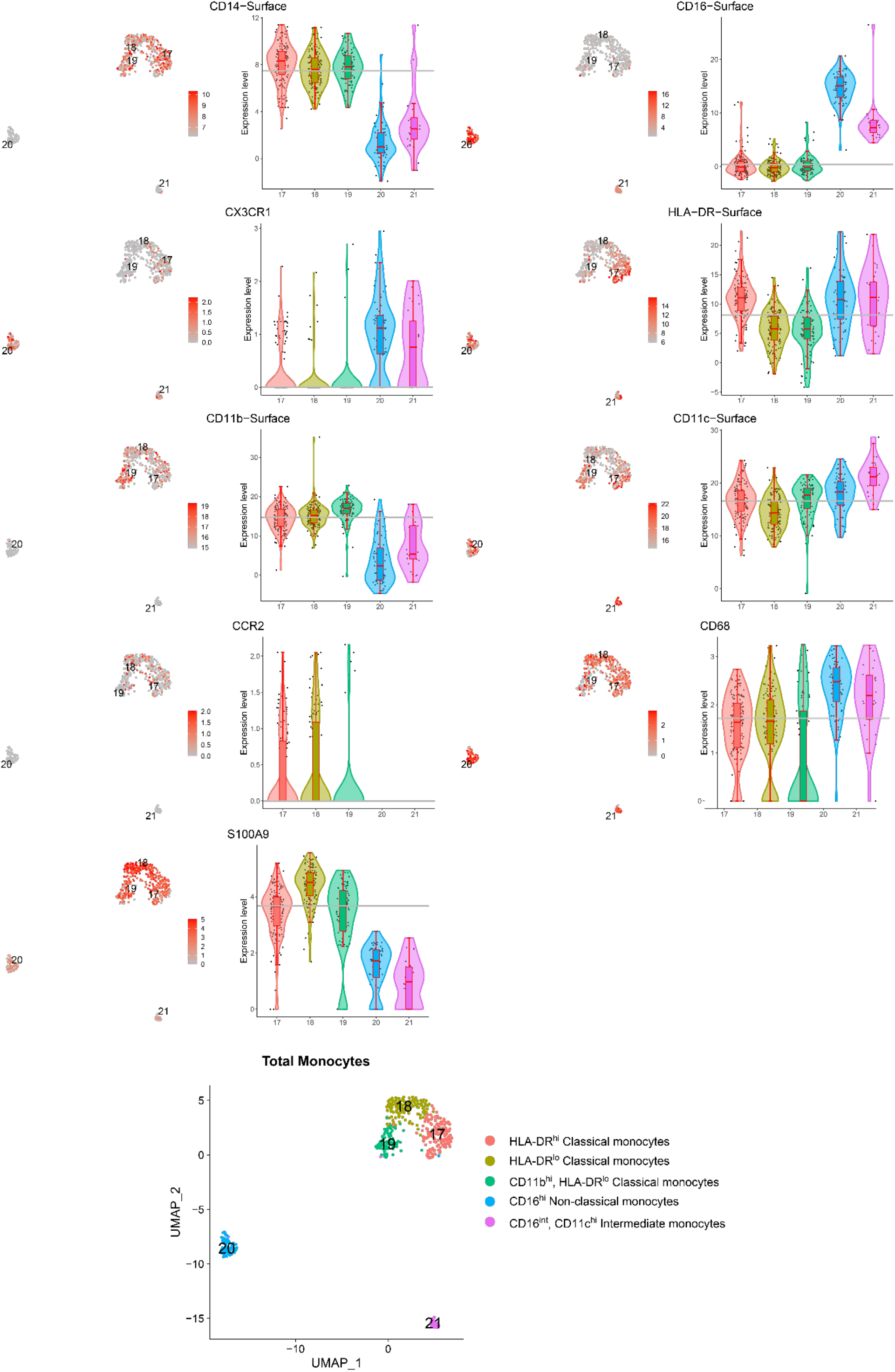
Blood monocytes identified by the SuPERR-seq workflow. UMAP and violin plots show cell-surface markers (ADT) and gene expression (GEX) levels used to identify and classify five subsets of monocytes in peripheral blood mononuclear cells (PBMC). The grey line shows the mean expression level across all clusters.

**Figure S7.**
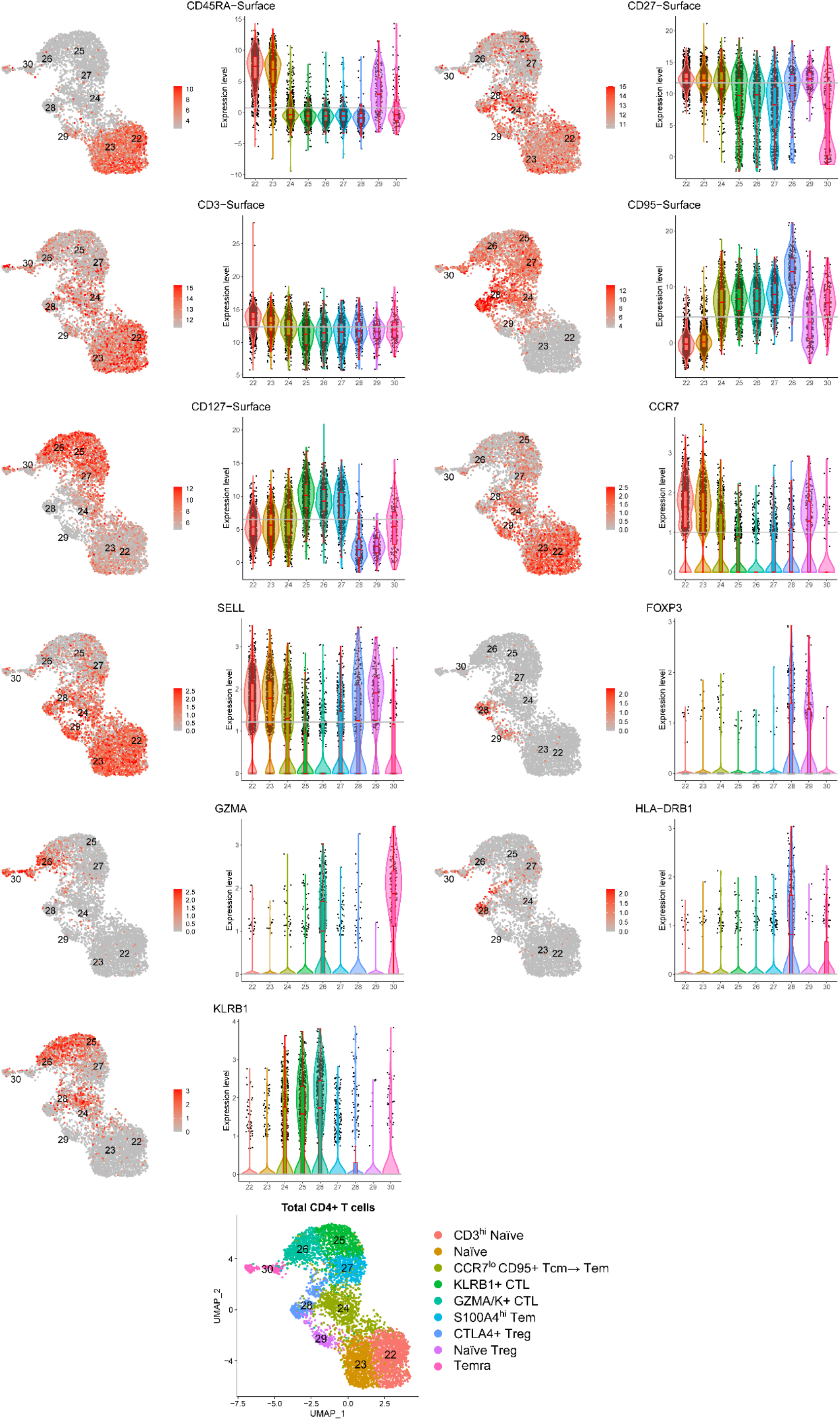
Blood CD4 T-cell subsets identified by the SuPERR-seq workflow. UMAP and violin plots show cell-surface markers (ADT) and gene expression (GEX) levels used to identify and classify nine subsets of CD4 T cells in peripheral blood mononuclear cells (PBMC). The grey line shows the mean expression level across all clusters. Tcm: central memory T cell; Tem: effector memory T cell; CTL: cytotoxic T lymphocyte; Treg: regulatory T cells; Temra: effector memory expressing CD45RA T cell.

**Figure S8.**
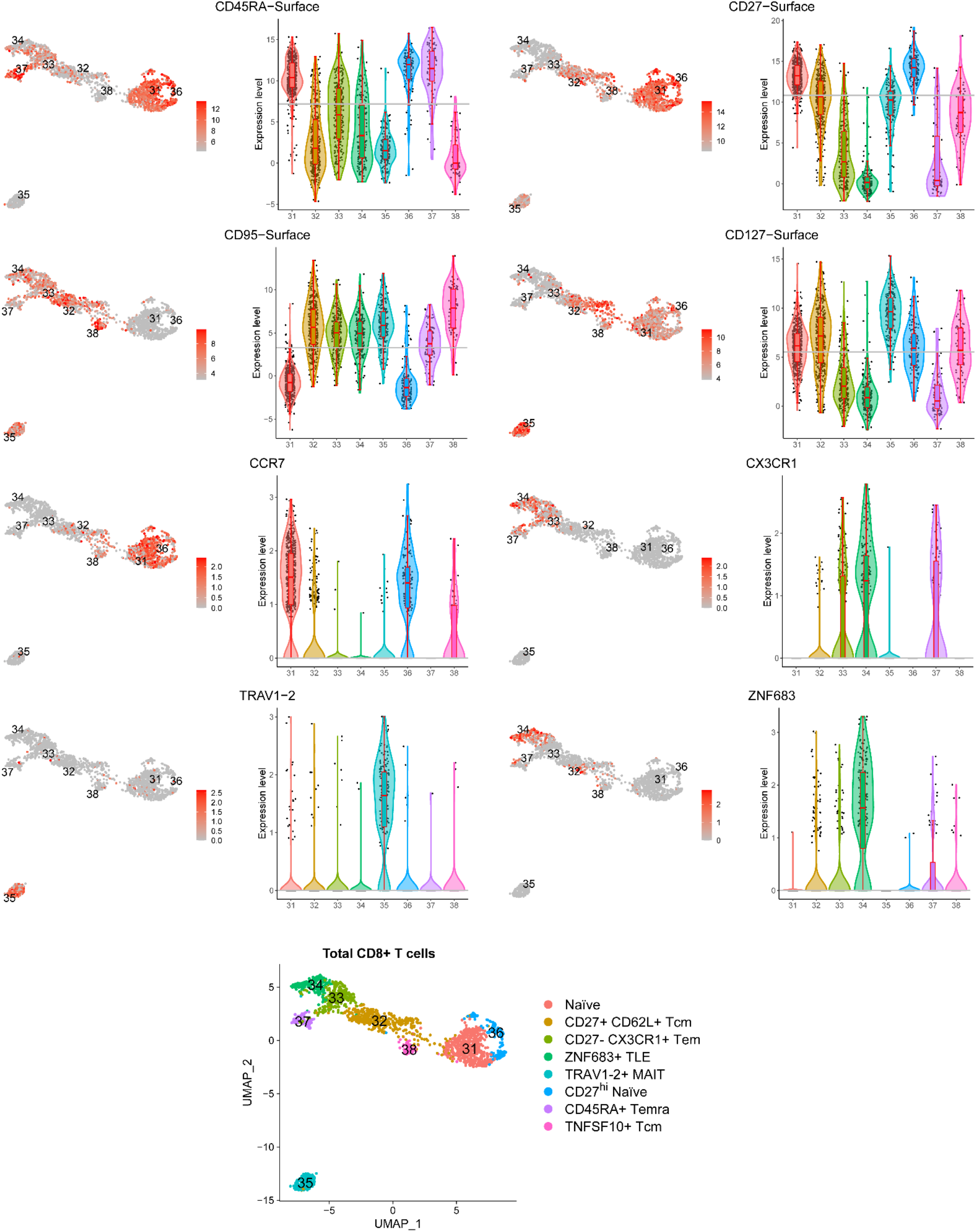
Blood CD8 T-cell subsets identified by the SuPERR-seq workflow. UMAP and violin plots show cell-surface markers (ADT) and gene expression (GEX) levels used to identify and classify eight subsets of CD8 T cells in peripheral blood mononuclear cells (PBMC). The grey line shows the mean expression level across all clusters. Tcm: central memory T cell; Tem: effector memory T cell; TLE: long-lived effector memory T cell; MAIT: mucosal-associated invariant T cells; Temra: effector memory expressing CD45RA T cell.

**Figure S9.**
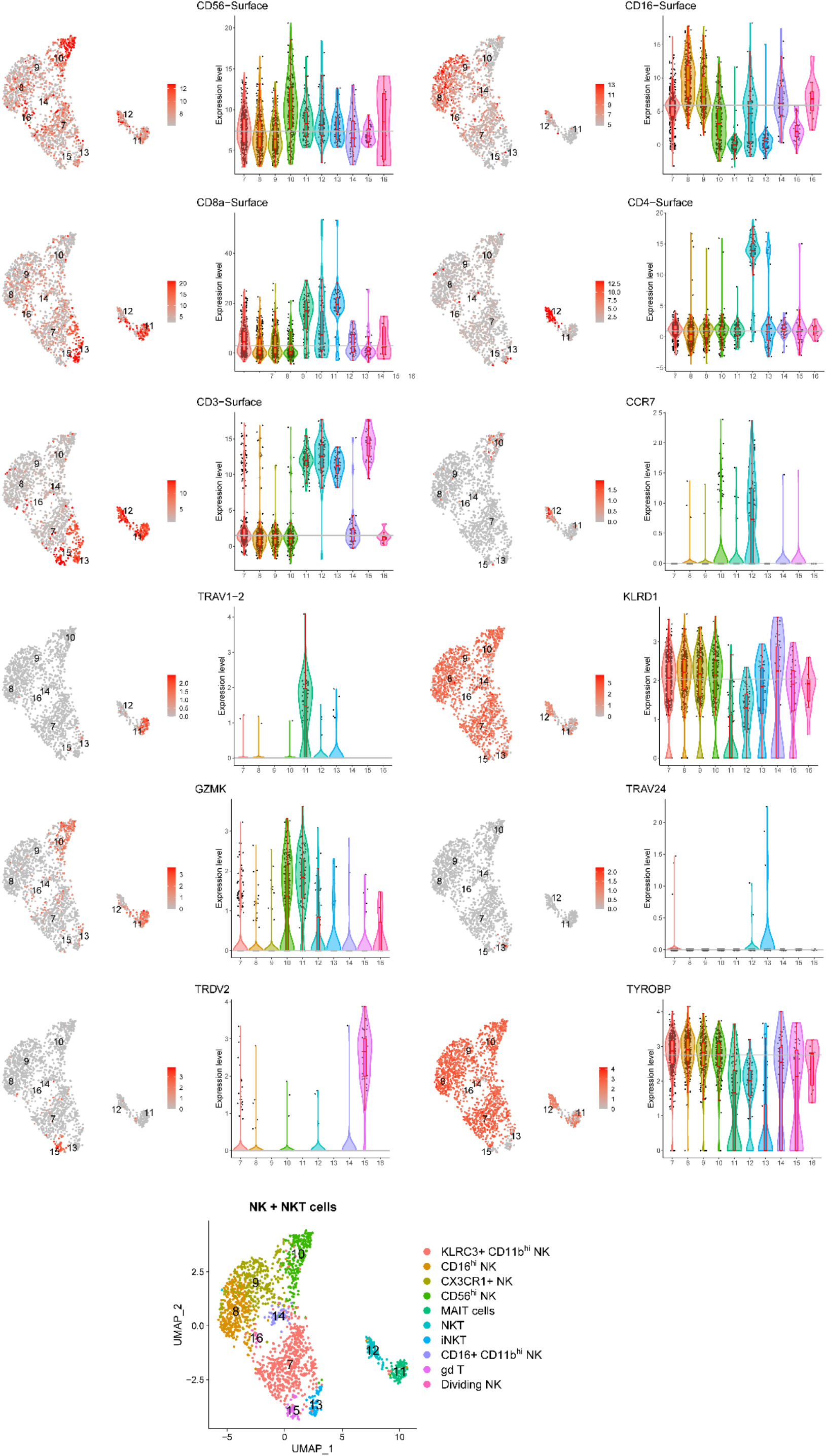
Blood NK, NKT, MAIT and γδ T cell subsets identified by the SuPERR-seq workflow. UMAP and violin plots show cell-surface markers (ADT) and gene expression (GEX) levels used to identify and classify ten subsets of NK, NKT, MAIT, and γδ T cells in peripheral blood mononuclear cells (PBMC). The grey line shows the mean expression level across all clusters. MAIT: mucosal-associated invariant T cells; NKT: natural killer T cells; iNKT: invariant NKT; γδ T: gamma-delta T cells.

**Figure S10.**
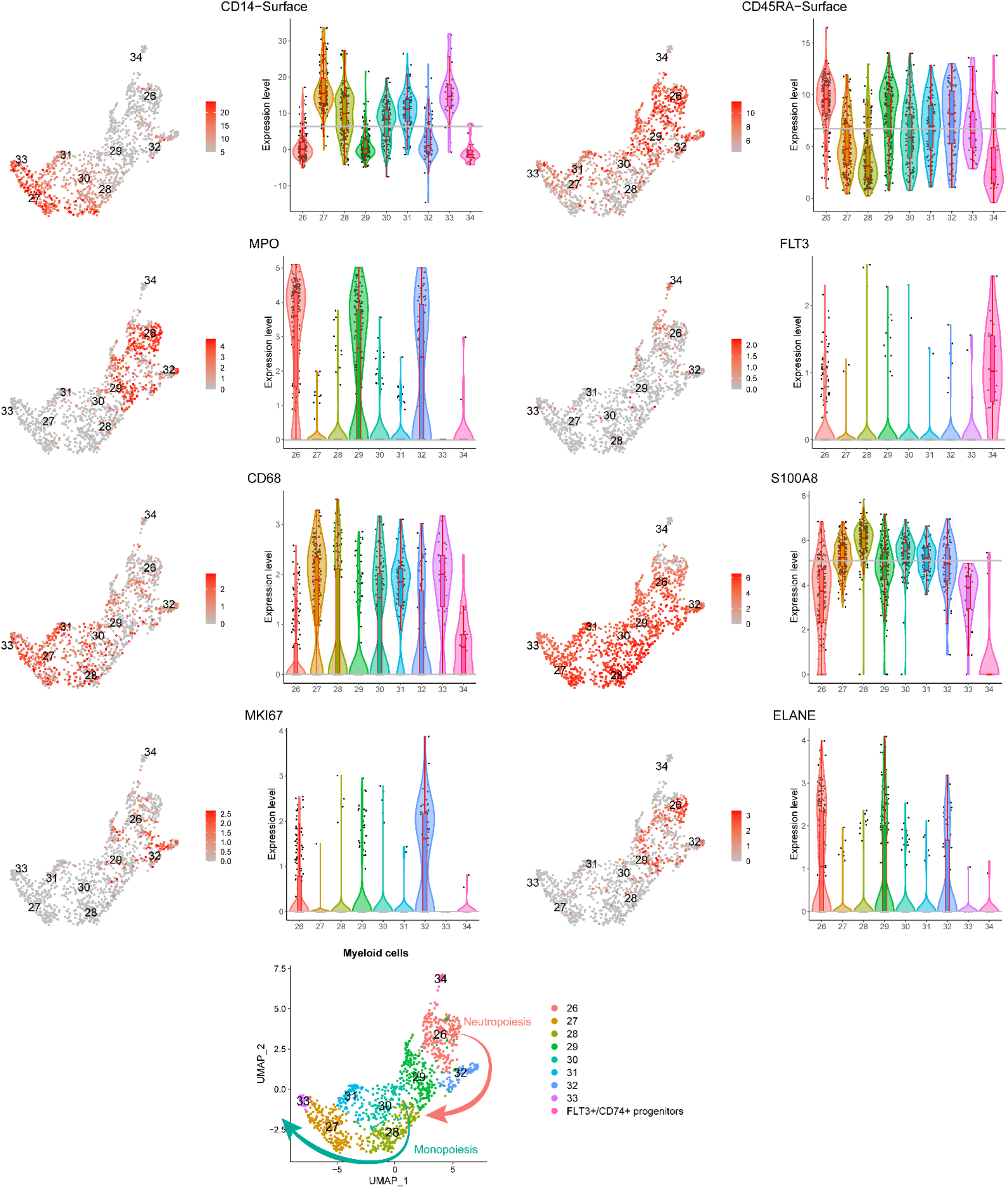
Monocyte and neutrophil development in the BM identified by the SuPERR-seq workflow. UMAP and violin plots show cell-surface markers (ADT) and gene expression (GEX) levels used to identify and classify nine subsets along the monopoiesis and granulopoiesis pathways in the bone marrow (BM). The grey line shows the mean expression level across all clusters. The red arrow shows the potential direction of the developmental pathway for neutrophils. The cyan arrow shows the potential direction of the developmental pathway for monocytes.

**Figure S11.**
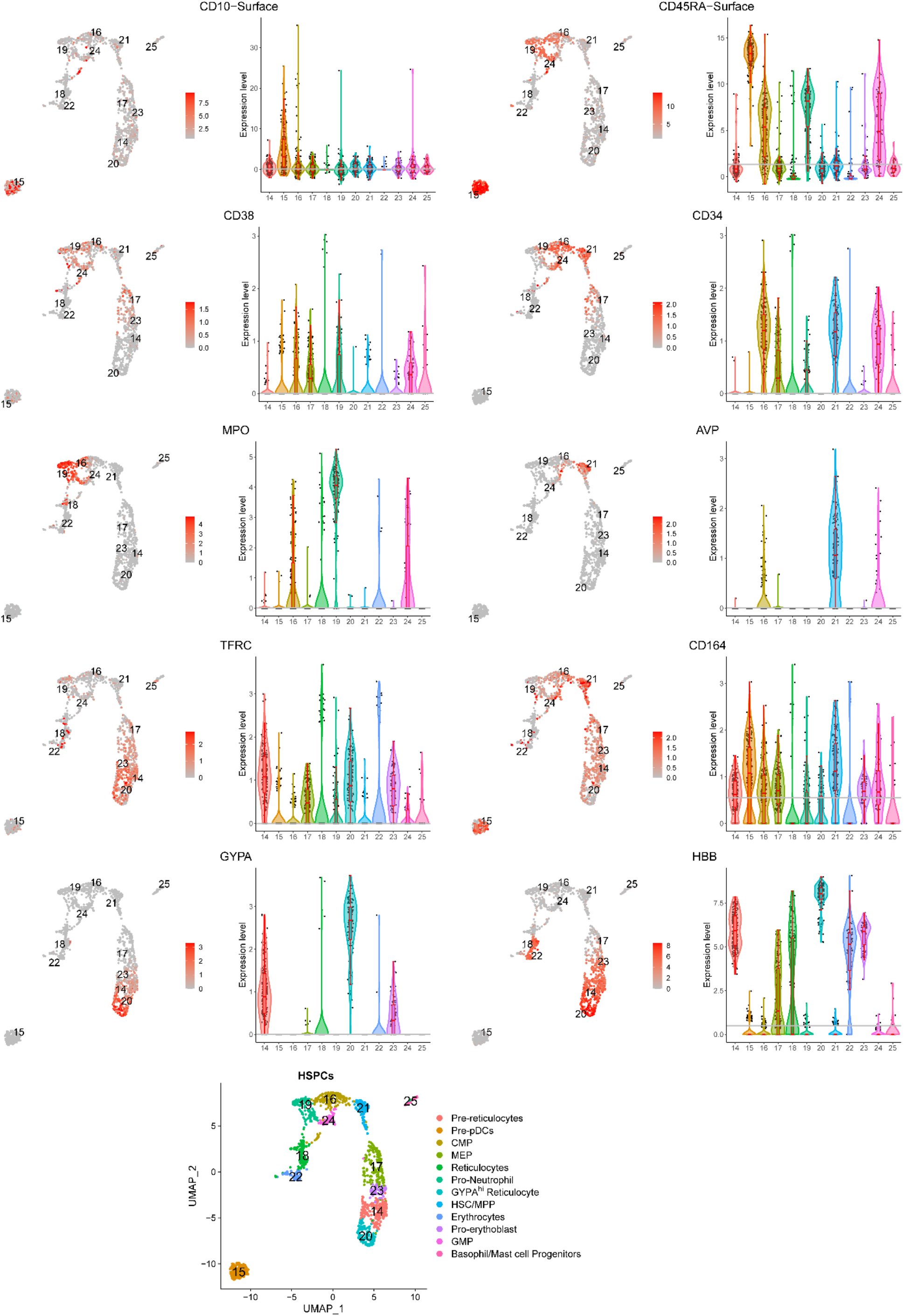
BM-derived Hematopoietic Stem and Progenitor Cells (HSPCs) identified by the SuPERR-seq workflow. UMAP and violin plots show cell-surface markers (ADT) and gene expression (GEX) levels used to identify and classify twelve subsets of HSPCs in the bone marrow (BM). The grey line shows the mean expression level across all clusters. Pre-pDCs: pre-plasmacytoid dendritic cells; GMP: granulocyte-monocyte progenitor; MEP: megakaryocyte-erythroid progenitor; HSC: hematopoietic stem cell; MPP: multipotent progenitor; CMP: common myeloid progenitor.

**Figure S12.**
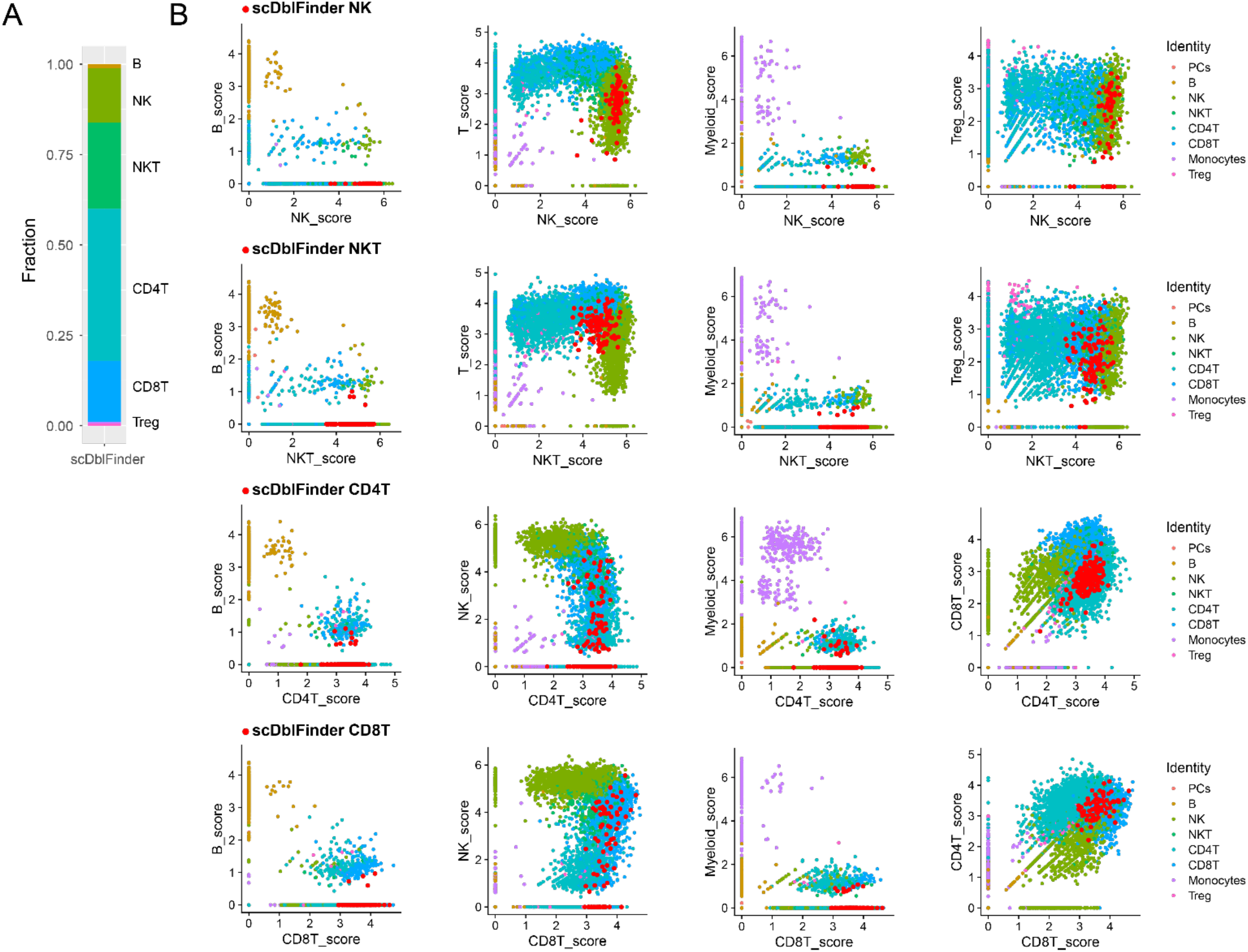
Gene expression profile of the heterotypic doublets identified by scDblFinder. (A) Proportion of heterotypic doublets identified in the peripheral blood mononuclear cells (PBMC). Cell types were annotated based on the SuPERR-seq workflow/classification. (B) Expression level of cell-type-specific gene signature/score (see Methods) in heterotypic doublets defined by scDblFinder (red points). Based on this gene expression analysis, we were not able to confirm whether the doublets defined by scDblFinder are indeed heterotypic doublets.

**Figure S13.**
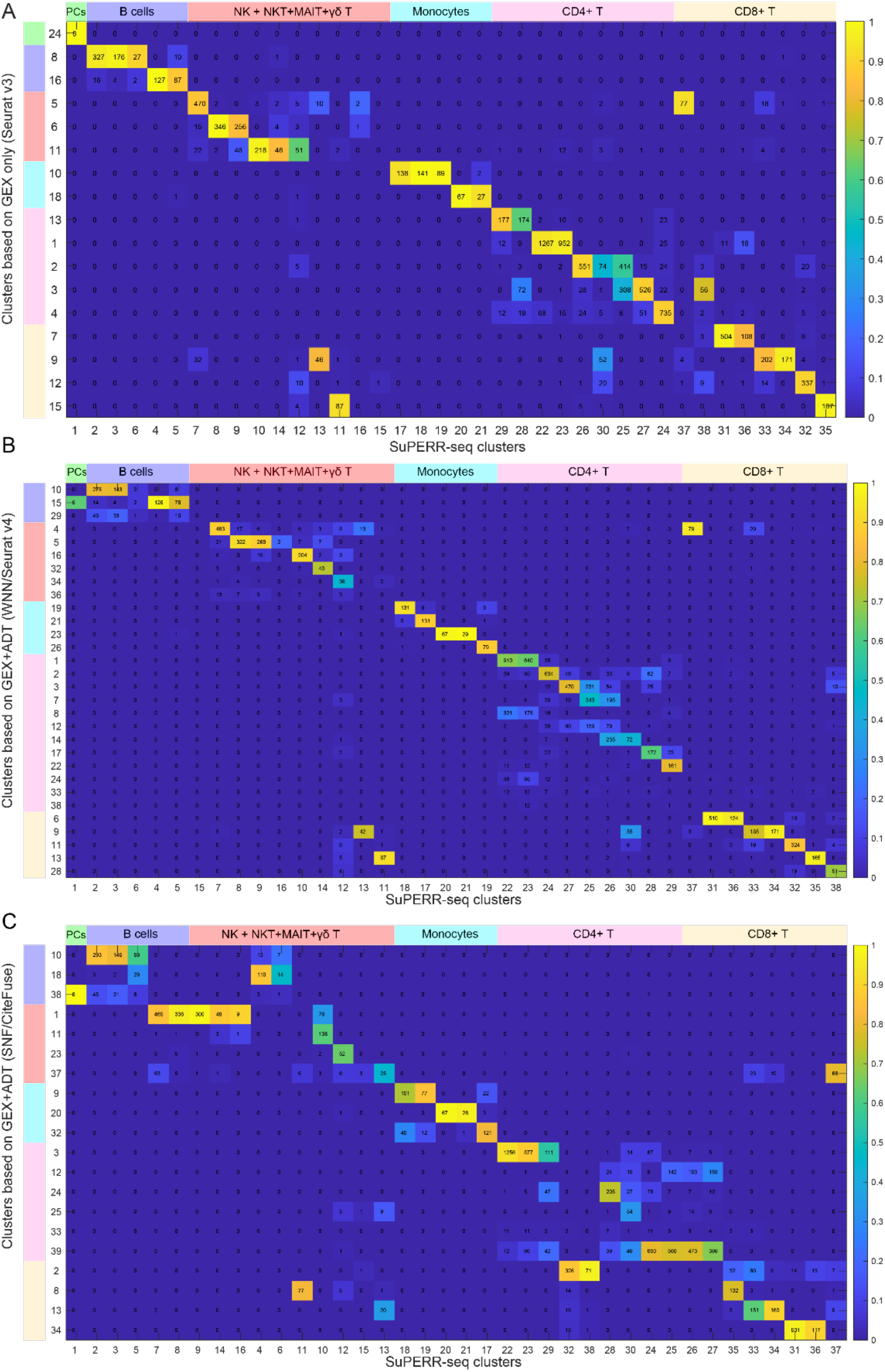
Comparisons between SuPERR-seq and other conventional approaches reveal cell-type misclassifications (peripheral blood). (A) Cross comparison of clustering results between SuPERR-seq and Seurat v3. (B) Cross comparison of clustering results between SuPERR-seq and WNN/Seurat v4. (C) Cross comparison of clustering results between SuPERR-seq and SNF/CiteFuse. For the sake of comparison, we ran each workflow using the same set of cell barcodes. None of the workflows was able to identify the plasma cells (PCs) cluster that was defined by the SuPERR-seq “manual gating” step using the ADT and Ig-specific transcripts (see Fig. 7A). The PCs were included here as SuPERR-seq cluster 1.

**Figure S14.**
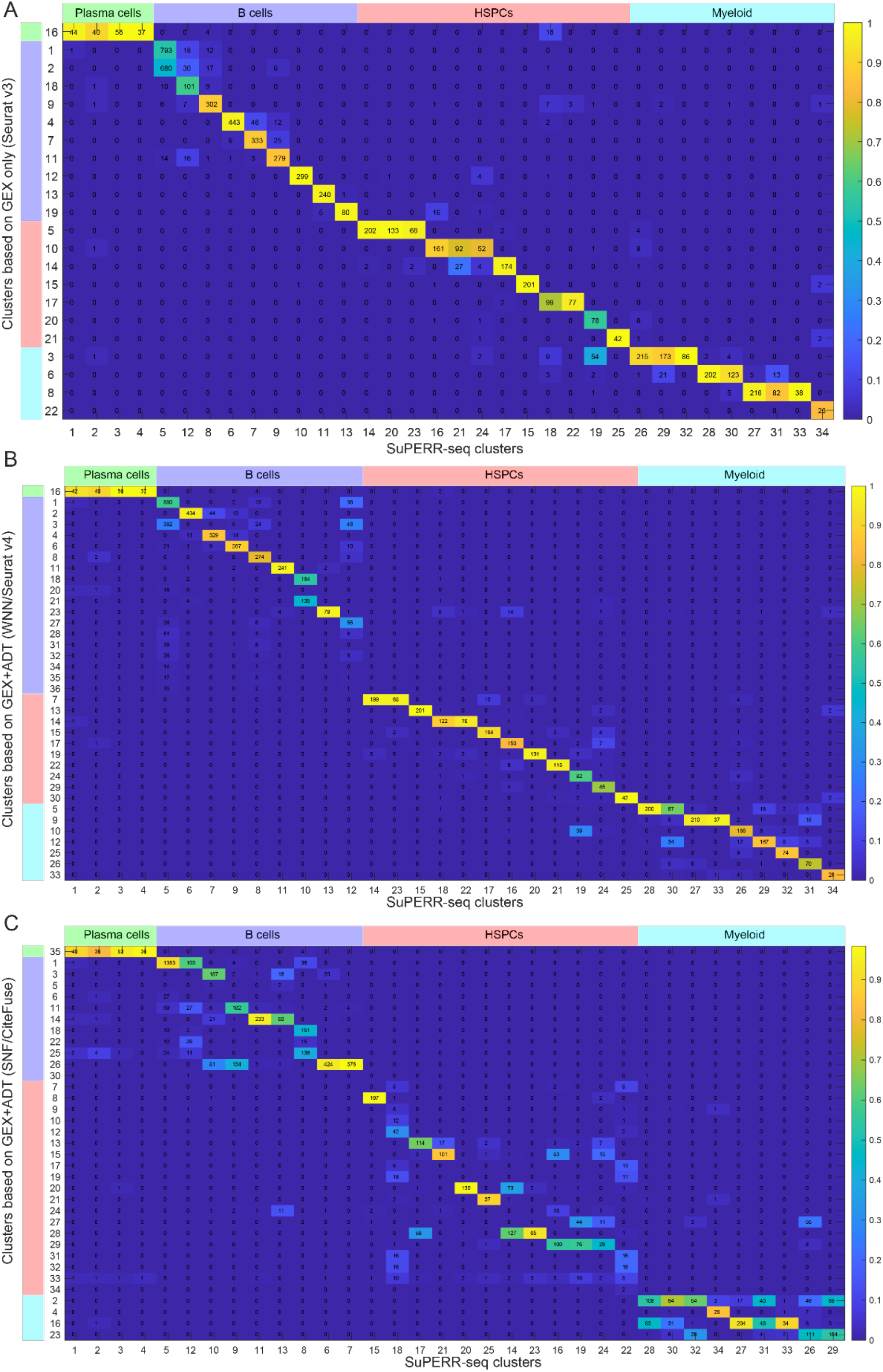
Comparisons between SuPERR-seq and other conventional approaches reveal cell-type misclassifications (bone marrow). (A) Cross comparison of clustering results between SuPERR-seq and Seurat v3. (B) Cross comparison of clustering results between SuPERR-seq and WNN/Seurat v4. (C) Cross comparison of clustering results between SuPERR-seq and SNF/CiteFuse. For the sake of comparison, we ran each workflow using the same set of cell barcodes.

**Figure S15.**
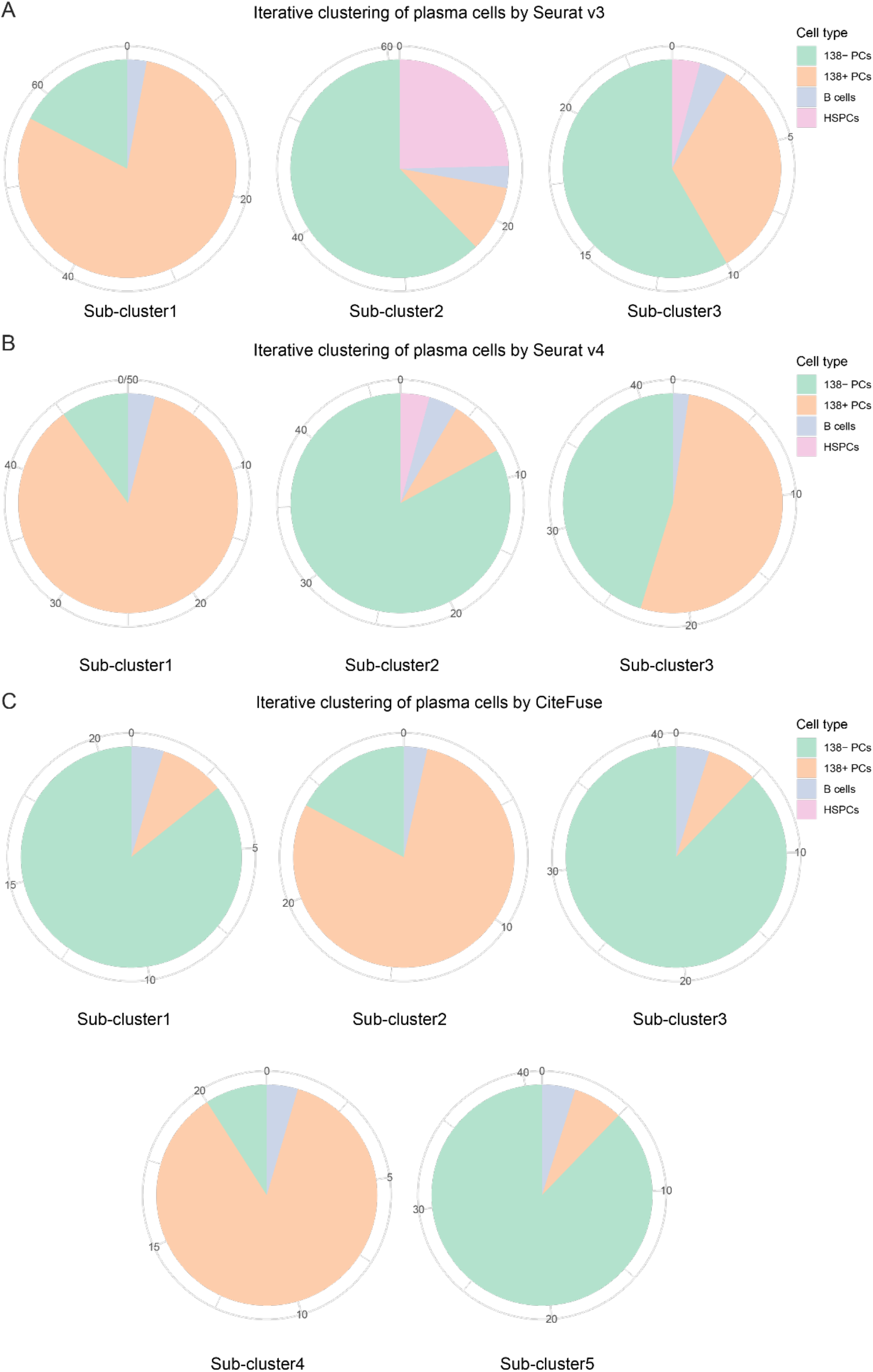
Sub-clustering of the bone marrow plasma cell cluster identified by the conventional workflows generated misclassified subsets. (A) Louvain sub-clustering of plasma cells (PCs) defined by Seurat v3. (B) Louvain sub-clustering of PCs defined by Seurat v4. (C) Louvain sub-clustering of PCs defined by CiteFuse.

